# Microenvironmental factors in cell segregation and heterogeneity in breast cancer development

**DOI:** 10.1101/2021.12.01.470838

**Authors:** J. Roberto Romero-Arias, Carlos A. González-Castro, Guillermo Ramírez-Santiago

**Affiliations:** Instituto de Investigaciones en Matemáticas Aplicadas y en Sistemas, Universidad Nacional Autónoma de México, 01000 Ciudad de México, Mexico; Instituto de Matemáticas, Universidad Nacional Autónoma de México, Juriquilla Querétaro, Mexico

**Keywords:** Microenvironmental agents, Estrogens, Heterogeneity, Cancer mutations

## Abstract

We analyzed a quantitative model that describes the epigenetic dynamics during the growth and evolution of an avascular tumor. A gene regulatory network (GRN) formed by a set of ten genes that are believed to play an important role in breast cancer development was kinetically coupled to the microenvironmental agents: glucose, estrogens and oxygen. The dynamics of spontaneous mutations was described by a Yule-Furry master equation whose solution represents the probability that a given cell in the tissue undergoes a certain number of mutations at a given time. We assumed that mutations rate is modified by nutrients spatial gradients. The tumor mass was grown by means of a cellular automata supplemented with a set of reaction diffusion equations that described the transport of the microenvironmental agents. By analyzing the epigenetic states space described by the GRN dynamics, we found three attractors that were identified with the cellular epigenetic states: normal, precancer and cancer. For two-dimensional (2D) and three-dimensional (3D) tumors we calculated the spatial distributions of the following quantities: (i) number of mutations, (ii) mutations of each gene and, (iii) phenotypes. Using estrogens as the principal microenvironmental agent that regulates cells proliferation process, we obtained the tumor shapes for different values of the estrogen consumption and supply rates. It was found that he majority of mutations occurred in cells that were located close to the 2D tumor perimeter or close to the 3D tumor surface. Also It was found that the occurrence of different phenotypes in the tumor are controlled by the levels of estrogen concentration since they can change the individual cell threshold and gene expression levels. All the results were consistently observed for 2D and 3D tumors.

## INTRODUCTION

In the past decades the number and variety of quantitative models on cancer has increased significantly. They have been aimed at addressing important aspects of cancer development such as tumor initiation and progression, tumor-structure, intra-tumor-heterogeneity as well as genetic evolution (1–6). Nonetheless, the high complexity of cancer posses challenges and new opportunities for novel approaches and more elaborated quantitative models. Quantitative modeling has the ability to reveal unknown and/or unexpected biological as well as physical features and predictions that can be validated experimentally and clinically. Currently there is no consensus over how cancer is initiated, however, it is known that tumor growth happens in several different stages. The general accepted view is that a cell must undergo series of gene mutations before it becomes cancerous. That is, cancer development is the result of the gradual accumulation of mutations that enhance cell proliferation rate and inhibit cell death rate leading to tumor progression (7–9). The detailed factors that drive these mutations are unknown, nonetheless, there is a general credence that environment and heredity play important roles in cancer initiation. The external environmental effects upon genes can influence dramatically cells behavior, and some of these effects can be inherited.

Epigenetics studies heritable phenotype changes that do not involve alterations in the DNA sequence. That is, epigenetic changes can influence gene expression without a change in genotype, and determine which proteins are transcribed. Genotype describes the complete set of genes in an organism. While different genotypes can give rise to different phenotypes, the microenvironment in which the cell or organism develops can also change the expressed phenotype. It has been observed that even genetically identical individuals growing in the same microenvironment can be very different (10).

The phenotypic plasticity of microorganisms allows them to maintain optimal growth rates even under adverse conditions. Some studies suggest that cells can express metabolic enzymes at tight concentrations by adjusting their gene expression. In fact, the associated transcription factors are often regulated by intracellular metabolites. In single-celled organisms, regulatory networks respond to the microenvironment optimizing processes for their survival. In multicellular animals, the same principle has been put at the service of the cascades of genes that control the shape of the organism. Every time a cell divides, it gives rise to two cells, however, they may differ in the genes that are activated in spite of the fact that their complete genome is the same. Often times a “self-sustaining feedback loop” ensures that a cell maintains its identity and transmits it to its descendants.

The mechanisms that integrate signal transduction and cell metabolism are largely conserved between normal and cancer cells. Nonetheless, normal cells require an structured mechanism, that involves extracellular stimulation, growth factors and downstream signaling pathways, for proliferation and conserved gene expression and cell physiology. Meanwhile, cancer cells can increase metabolic autonomy and often times undergo mutations that chronically enhance these pathways, allowing them to maintain a metabolic phenotype of biosynthesis independently of normal physiologic constraints. A more complete understanding of the metabolic phenotype of cell proliferation is still to be discovered. Under this view, one of the main goals of future research is to determine the impact of signaling mediators on specific and global metabolic activities.

In breast cancer one can consider that the most important epigenetic factors are metabolism (11–16) and estrogen production (17–21). The former contributes to reprogramming several metabolic pathways that are essential for cancer cell survival and tumor growth while the latter affects significantly the cell proliferation process. However, there are other features that allow tumor cells to take up abundant nutrients and use them to produce ATP, generate biosynthetic precursors, and tolerate stresses associated with malignancy, for instance, redox stress and hypoxia. In addition, there is an emerging class of reprogrammed pathways that involve those allowing cancer cells to tolerate lack of nutrients by catabolizing macromolecules from inside or outside the cell. For example, autophagy, macropinocytosis, and lipid scavenging. This reprogramming may be regulated intrinsically by tumorigenic mutations in cancer cells or extrinsically by influence of the microenvironment (16). On the other hand, estrogens play a major role in promoting the proliferation of both, normal and neoplastic breast epithelium. However, there is no clear understanding of the mechanisms through which estrogens cause cancer (18). The most widely acknowledged mechanism of estrogen carcinogenicity is its binding to its specific nuclear receptor alpha (ER-α) for exerting a potent stimulus on breast cell proliferation through its direct and/or indirect actions on the enhanced production of growth factors (22–24).

The incorporation of epigenetics and estrogens production into a quantitative model of cancer evolution allows the integration of both, intrinsic (biochemical) and extrinsic (microenvironmental) signals in the genome dynamics during the development of breast cancer. This paper proposes and analyze a quantitative model that incorporates these two processes by considering some metabolic aspects and the role of estrogens during cell proliferation. The former are believed to be the biochemical substances that trigger many signaling pathways during breast cancer evolution while the latter promotes proliferation of normal and neoplastic cells. To the best of our knowledge, up to now, there is no quantitative model that considers such integration in breast cancer evolution.

## MODEL

The growth and development of an avascular tumor is analyzed by considering a cellular automata together with a set reaction-diffusion equations (25, 26) that describe the transport of essential nutrients: glucose, oxygen and estrogens. It is assumed that tissue is made of four types of cells, namely: normal, precancer, cancer, and tumor necrotic cells that live in either, a 2D square lattice or a 3D cubic lattice. Normal and necrotic cells may occupy one lattice site, however, more than one precancer or cancer cells can pile up at a given lattice site. According to this, three field variables are defined at each lattice site: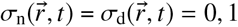, for normal and necrotic cells, and 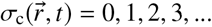 for precancer or cancer cells. The vector 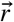 has two integer components, *i, j* in 2D or three integer components *i, j, k*, in 3D with 0≤ *i, j, k*≤ *L*. They denote the cell position coordinates in the lattice. An initial cancer cell is placed at about the middle of the lattice and a nutrient supply –horizontal capillary vessel– is located at the upper side. It was assumed that essential nutrients, oxygen and estrogens diffuse from the capillary vessel throughout the tissue.

Nutrients and estrogens are critical for DNA synthesis as well as for cell proliferation (22, 23, 27–29). Thus, it can be considered that nutrients play the role of catalysts during the expression of genes whereas estrogens regulate the cell cycle and different signaling pathways. Additionally, it is assumed that there exits a competition between normal and cancer cells for essential nutrients, oxygen and estrogens. The abundance of these substances around cells yields fluctuations and asymmetries in gene propensities, which in turn play a role in the development of heterogeneity. On the other hand, the growth of tumor cells is typically limited to a region of approximately 10 cells from a blood vessel that supplies the tumor. As a result, there appear nutrients, oxygen and estrogens spatial gradients (30). Taking this fact into consideration, the transport of nutrients, oxygen, and estrogens in the tissue is described by the following set of three reaction-diffusion equations (25, 26):

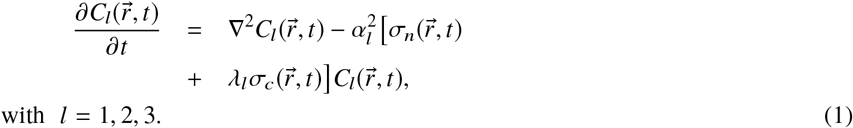

The quantities 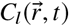, represent the glucose (*l* = 1), oxygen (*l* = 2) and estrogen (*l* = 3) concentration, respectively. The parameters 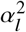 and λ_*l*_ represent the cells glucose (*l* = 1), oxygen (*l* = 2), and estrogens (*l* = 3), consumption and supply rates, respectively. The ability of the normal and cancer cells to compete for glucose, oxygen and estrogens are represented by the product, 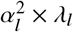. A transport equation for estrogens has also been included in Eqns. (1) since we have assumed that estrogens enhance the production of growth factors promoting cell proliferation of both, normal and neoplastic breast cells. The processes of cancer cell division and death are described independently in the next subsections by probability distributions that depend on nutrients, oxygen and estrogens local concentrations.

### Oxygen and death

Over the past decades, evidence has been accumulated showing that 50%-60% of advanced solid tumors develop hypoxic and/or anoxic tissue regions that are heterogeneously distributed within the tumor mass. Oxygen-sensing mechanisms have been developed in mammals to maintain cell and tissue homeostasis, as well as to adapt to the chronic low-oxygen conditions found in cancer. It involves the capture, binding, transport, and delivery of molecular oxygen. One of the crucial features of this network is its ability to sense and respond to low-oxygen concentration conditions. The poor vasculature in the early tumor development –avascular tumor– alters its metabolism (31). As a result there are too many tumor regions that undergo hypoxic stress because they are located at relatively large distances from blood vessels (32). Therefore, tumor cells have to adapt their metabolism to this unusual and harsh microenvironment that contains very small concentrations of nutrients and oxygen (33).

Under these conditions, hypoxia inducible factors (HIFs) activate for the maintenance of cellular oxygen homeostasis and hypoxia adaptation (34, 35). Oxygen is crucial in controlling vascularization, glucose metabolism, survival and tumour spread. This pleiotropic action is orchestrated by HIF, which is a master transcriptional factor in nutrient stress signalling (36). In a hypoxic environment accelerated glycolysis ensures ATP levels that are compatible with the demands of the fast proliferating tumor cells. This shift in cellular metabolism from mitochondrial respiration to glycolysis is linked to tumor malignancy. Sustained tumor hypoxia also gives rise to adaptations which allow cells to survive and even thrive. As a consequence, a more malignant phenotype may develop due to the following factors: (i) HIF-1*α*-mediated mechanisms favoring tumor growth and malignant progression, (ii) HIF-1*α*-independent up-regulation and down-regulation of genes, and (iii) effects via genome changes, that produce hypoxia-induced apoptosis resistance, genomics instability which in turn lead to clonal heterogeneity, and selection of resistant and/or aggressive clonal variants (37). On the other hand, it has been found that the rate of cell death increases when supply of glucose and nutrients are very low and cellular ATP is increasingly depleted. The most striking proof of hypoxia-induced apoptosis is the suppression of the electron transport chain on the inner membranes of the mitochondria (38). Taking these observations into consideration one can assume that cell death occurs with certain probability that depends on the local oxygen concentration as in (25, 26).

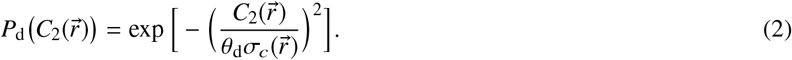

This probability is represented by a sigmoidal curve with parameter θ_d_ that controls its shape, 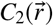 is the local oxygen concentration, and 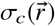 represents the number of cancer cells located at point 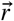. This approximation can also explore the fact that exposure to cycling hypoxia leads to high levels of reactive oxygen species (ROS) from cell mitochondria. On the othe hand, it has been demonstrated that reoxygenation induces significant amounts of DNA damage and malignant progression because of the presence of ROS during reoxygenation (39, 40). Thus, ROS influences the tumor microenvironment and is known to initiate cancer angiogenesis, metastasis, and survival at different concentrations (41). However, the detailed molecular mechanisms of how cancer cells respond to oxidative stress are still to be discovered.

### Estrogens and Cell Cycle

Estrogens give rise to diverse biological effects as a result of its direct interaction with an intracellular receptor that activates the expression of genes encoding proteins with important biological functions (42–45). In terms of the molecular mechanism of action, estrogens induce gene expression and synthesis of specific proteins, activation of specific enzymes, and proliferation in certain cell types. All of these actions appear to require the binding of the hormone to a specific receptor protein (46). In particular, one of the most notable estrogens effect is its potential mitogenic action in hormone sensitive breast epithelial tissues (47, 48). Because of this, the establishment of estrogens concentration gradients are crucial for cell proliferation and cancer progression. By contrast, nutrients availability facilitates nutrients cell consumption which favors mutation rates whereas limited or lack of nutrients lead to latent cell states with small mutation rates. This latter hypothesis was modeled as a stochastic process which was coupled to the nutrients transport reaction-diffusion equations. The dynamics described with this coupling yielded genetic spatial heterogeneity as a result of mutations accumulation during tumor growth, a hallmark of most cancers (26).

Clinical and animal studies suggest that risk factors associated with breast cancer reflect cumulative exposure of the breast epithelium to estrogens (49). During the cell cycle estrogens regulate the expression and function of cyclins, c-Myc, cyclin D1 and cyclin E-Cdk2 which are considered important in the control of G1/S phase progression (50, 51). Furthermore, Cyclin E shows an expression periodic pattern, being synthesized during the G1-phase of the cell cycle, with sharp increasing levels during the late G1 phase, followed by the accumulation of cyclin E protein and then down regulates in the S-phase. Other CDKs complexes, as cyclins A and B, undergo opposite periodic patterns to those cyclin E undergoes (51). Taking all the previous statements into consideration one can surmise that the period of the cell division cycle is mainly regulated by the local concentration of estrogens 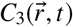.

This antagonistic process leads to out–of phase non-linear oscillations with maxima related to the transitions of the cell cycle G phases, changing the entire cell cycle (52). Bearing this in mind, we assumed that cyclins E and B are the two key players responsible for the out of phase oscillations. To describe these out of phase –opposite– periodic oscillation patterns of the relative concentrations of cyclins B and E that drive the cell cycle we have chosen the following relatively simple non-dimensional Lotka-Volterra system of equations.

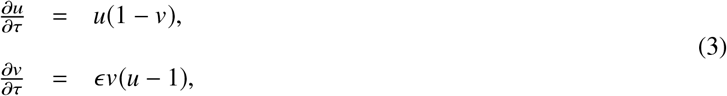

where *u* and *v* represent the concentrations of cyclins E and B, of a cell located at point 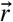 at time τ, respectively. The wave shape solution depends only on local parameter 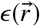 and on the boundary conditions for each cell. This system describes out of phase oscillations with period 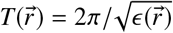.

Experimental data has shown that the cell cycle is arrested if the estrogen concentration is below or above certain threshold value (53). This fact allows one to assume that the growth rate depends on estrogen consumption rate. This quantity is directly related to the reaction term in the third of Eqns. 1, the reaction-diffusion equation for estrogens. For that reason the time scale of the oscillations can be related to the estrogens local concentration through the local parameter 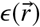 which can be expressed as

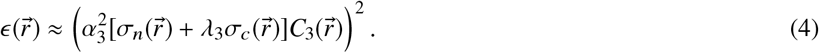

In principle, one can assume that normal and cancer cells follow a similar synchrony during cell cycle. Notwithstanding, cell proliferation rate depends on their own estrogen consumption rates. Therefore, as estrogens consumption rate increases, cancer cells growth factor increases as well.

### Cell lineage and microenvironment

A single cell cannot influence significantly its immediate surroundings. Nonetheless, through the expression of cooperative phenotypes a relatively large number of cells can change significantly their microenvironment (54). Tumors are made of multiple sub-populations of cells with different phenotypes, some of them are able to self renew, seed, maintain tumors, and provide a reservoir of resistant cells (55). In this way many cell phenotypes contribute to the growth and division processes of nearby cells by changing their local environment (53).

In cancer there are two major types of phenotypic plasticity: initiating and maintaining plasticity. Initiating plasticity is generated by the influence of the cell of origin and the specific driver mutations that occur during tumor formation. Maintaining plasticity is the result of genetic evolution and hierarchical and plastic inter-conversion between cellular phenotypes. These two forces collaborate to generate the tumor phenotypes that are diverse even within the same tissue (56). Furthermore, cellular mechanisms that govern lineage proliferation and survival during development might also underlie tumorigenic mechanisms.

Somatic genetic alterations show lineage-restricted patterns across human tumors, which indicates that genetic changes in cancer might be conditioned by the lineage programs embedded in tumor precursor cells (57). The preferential interaction among genetically related individuals increases the propensity for cooperative phenotypes to evolve. Different cell lineages segregate in space with the aim at cooperating and benefit each other. Because of this, local populations of cancer cells often times aggregate in groups of progenitors that proliferate leading to large clusters with similar lineages (58).

Here, we apply the concept of lineage cell segregation –that has been applied recently to analyze the evolution of bacterial colonies (54)– as a measure of cell heterogeneity during tumor development. Considering that estrogens induce gene expression, synthesize specific proteins, and activate specific enzymes which require the binding of the hormone to a specific receptor protein, one can model the lineage production rate with a metabolic Michaeles-Menten kinetics. In doing so, the segregation index at position 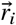 can be expressed as,

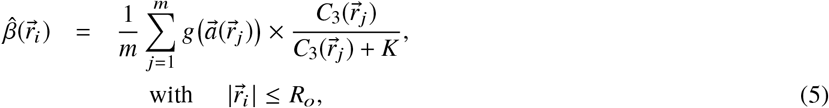

where 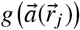 is the genetic activity defined by

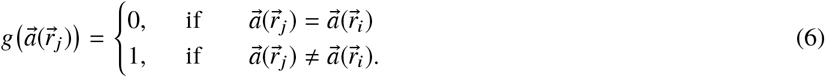

and 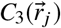 is the local estrogens concentration, *K* is the half saturation constant at the neighboring cells with positions 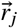 and *j* = 1, 2, 3,, *m*, is the total number of neighboring cells within a given distance *R*_*o*_ located at positions 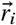. Note that the genetic activity discourages the interaction between cells of the same genotype because they do not lead to genetic diversity. The vector 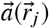 represents the genotype of cell located at position 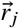 and is formed by the set of breast cancer genes that are believed to play a major role in cancer development. The gene set that completes the entries of vector 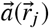 will be discussed in detail in subsection. The form of the segregation index directly measures the spatial assortment of genetic cell lineages and ranges from 0 to 1, where 1 denotes complete lineage segregation within the spatial scale *R*_*o*_.

### Random mutation dynamics

Cancer cell genome instability leads to a transient or persistent state that increases the spontaneous mutation rate that leads to a cascade of mutations some of which enable cancer cells to bypass the regulatory processes that control cell location, division, expression, adaptation and death (59). Mutations either arise from copying not repaired DNA damage or from errors that happened during DNA synthesis (60, 61). In order to describe quantitatively the cascade of spontaneous mutations, we assumed that they depend only on the previous genetic state. Thus, we modeled the mutations dynamics by means of a Yule-Furry Markovian process which is described by the following master equation (62).

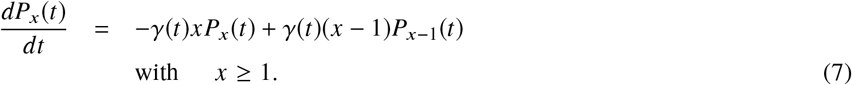

Here *P*_*x*_(*t*) represents the probability that a given cell in the tissue undergoes, *x* (*x* = 0, 1, 2,) mutations at a given time *t* with a hopping probability γ(*t*) > 0 that considers that one new mutation, *x* → *x* + 1, will happen in the time interval [*t, t* + *dt*). The solution of Eqn. (7) is a geometric probability distribution with argument, 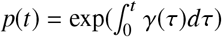 (62).

It has been found that cancer cells remodel tissue microenvironment and specialized niches to their competitive advantage (63, 64). Here we assume that nutrients spatial gradients change somehow the tissue microenvironment and cell niches modifying the acquisition rate of new spontaneous mutations (65, 66). Thus, one can surmise a direct relationship between nutrients Concentration and the hopping prob ability γ (*t*) of acquiring new mutations. Therefore, one can propose the following *anzats*,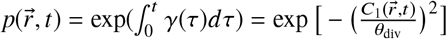, where 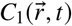 represents the nutrients concentration at the cell position 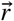 at time *t*, and θ_div_ is an adjustable parameter that controls the shape of the sigmoidal curve. Thus, there is an intrinsic nonlinear coupling between the master equation that describes the mutation dynamics of a given cell located at position 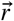 and the reaction-diffusion equation that describes the nutrients concentration at that location.

On the other hand, spontaneous mutations can be randomly activated by one of the following mechanisms: structural alterations resulting from mutation or gene fusion, by juxtaposition to enhancer elements, or by amplification of random mutations acquisition. These random activation can be modeled with a Poisson probability distribution (27, 67–69). Thus, the total probability distribution of having a mutation, at time *t*, at a given cell in the tissue is written as the product:

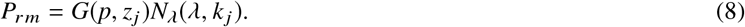

Here, *G*(*p, z*_*j*_) is a geometric probability distribution with mean (1 − *p*)/*p*, where *p* is given by the *anzats* indicated above, and *z*_*j*_ is the number of viable mutations of gene *j*. The Poisson probability distribution *N*_*λ*_(λ, *k* _*j*_) with mean λ represents the probability of occurrence of *k* _*j*_ mutations of gene *j* at a given cell. This factorization has proven to be very useful in the implementation of the stochastic simulations that describe tumor gene dynamics (26).

### Mutations induced by microenvironmental factors

There is a growing evidence suggesting that tumor microenvironment per se constitutes a significant source of genetic instability. The induction of mutagenesis and numerous types of DNA damage, including DNA strand breaks and oxidative base damage are associated with the adverse and harsh conditions of the microenvironment (65, 70). In this way, many solid tumors, including breast cancer, are composed of heterogeneous cell lineages that interact through complex networks and microenvironment (71). As in developing organs, tumor cells interact with these diverse cellular lineages, and regulate the hierarchy of tumorigenic cells (72). According to this view the local segregation index, 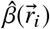, can be thought of as a probabilistic measure of the occurrence of mutations as a result of genetic instability induced by metabolic and micro-environmental conditions.

In the following paragraphs we consider the influence of nutrients and estrogens on mutations. According to the above proposed *anzats*, the probability *P* (*A*) = *p*, in the master Eqn. (7), represents the probability of occurrence of genetic mutations as a result of nutrients concentration gradients. On the other hand, most of the estrogens actions on the normal and neoplastic mammary cells are mediated via estrogen receptors, mainly for controlling cell proliferation. Failure of estrogen receptors increases the probability of mutations, *P* (*B*), due to the occurrence of genotoxic estrogen metabolites driven by estrogen concentration gradients (73).

Because of this the probability *P* (*B*) should be accounted in the mutation dynamics, however, its mathematical expression is unknown and it is very difficult to estimate. In spite of this limitation, here we explore its role by considering it as a varying parameter. Bearing this in mind, the probability that both, nutrients and estrogen concentration gradients influence mutations simultaneously but independently can be written as:

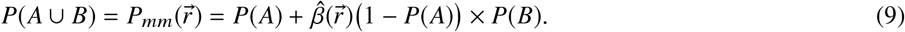

In this equation the segregation index 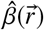 was introduced as a factor of *P* (*B*) because estrogens control cell proliferation and they can be considered as the main source of cell lineages. The segregation index measures the degree to which co-localized, metabolically active cells are genetically related to each other. Notice that when estrogens spatial gradients become negligible, *P*(*B*) = 0, then mutations are directly related to nutrients spatial gradients as has been analyzed in a previous model (26).

### Gene Regulatory Network

Gene Regulatory Networks (GRN) are an effective and useful models that are commonly used to study the complex regulatory mechanisms of a cell. A GRN is the collection of molecular species and their interactions which control gene-product abundance (74). GRN encode the patterns of interacting signals responsible for the up and down regulation of genes. GRNs integrate internal and external signals to ensure that a cell exhibits a response appropriate to its current environment (75) and moves around as the expression pattern changes (76).

A GRN evolves in time due to the mutual regulation of the genes, its dynamics eventually settles down into an equilibrium state that complies with the regulatory interactions (77). Here, the most notable topological features of a GRN state space are its basins of attraction. A basin of attraction in the state space is the set of states that moves over time toward a particular region called an attractor. An attractor corresponds to the steady states (or stable oscillations) of the system, while the a basin of attraction is formed with the set of initial system states that converge to a particular steady state. Transitions between attractors are triggered by regulatory signals that change the expression status of a set of genes in a concerted manner or by gene expression noise which produces random fluctuations in the expression of the genes (77). Because of this, attractors encode specific genetic programs of the cell that are already pre-programmed in the GRN, including those which produce a stable cell type-specific gene expression pattern. The detailed states space topography must be such as to guide the coordinated production of the appropriate proportions of distinct cell types at the right place and time, leading to the epigenetic landscape. Understanding the dynamics of GRN’s is crucial to comprehend development and the occurrence of diseases such as cancer.

The interplay between regulatory networks and metabolism and how an organism adapt to its microenvironment can also be described by GRNs (74, 78). In the epigenetic landscape not all attractors represent physiological cell phenotypes. The unused attractors are inevitable by products of the complex dynamics of the GRN, and the majority of them are associated to abnormal, non-viable gene expression patterns, that may be the result of conflicting signals. Therefore, cancer can be viewed as a set of abnormal gene expression patterns that correspond to attractors in the dynamics of an appropriate GRN. A first general description of this simple but profound idea was proposed fifty years ago by Kauffman (79).

#### Kinetic model

In this work, we consider a kinetic model of molecular regulatory interactions between a set of ten genes: TP53, ATM, HER2, BRCA1, AKT1, ATR, CHEK1, MDM2, CDK2 and P21 with transcriptional positive and negative regulations at the level of a single cancer cell (in appendix A is described the role each gene). It allows to uncover interesting macroscopic cancer phenomena at the physiological level which are observed in experiments and clinical trials, such as the phenotypic equilibrium in populations of breast cancer cell lines. In Fig. 1, we show the GRN composed by the genes indicated above together with their own regulatory links. Note that the genes related to some protein kinases, ATR, ATM, CHEK1, and CDK2 have a direct relationship with estrogens and the cell cycle. On the other hand, the BRCA1 and HER2 oncogenes can change their expression levels and modify the cell cycle when the concentration of estrogens in the environment changes. Additionally, tumor suppressor genes and oncogenes TP53, AKT1, MDM2 and P21 follow a relationship with oxidative stress or oxygen concentration in the environment.

**Figure 1:**
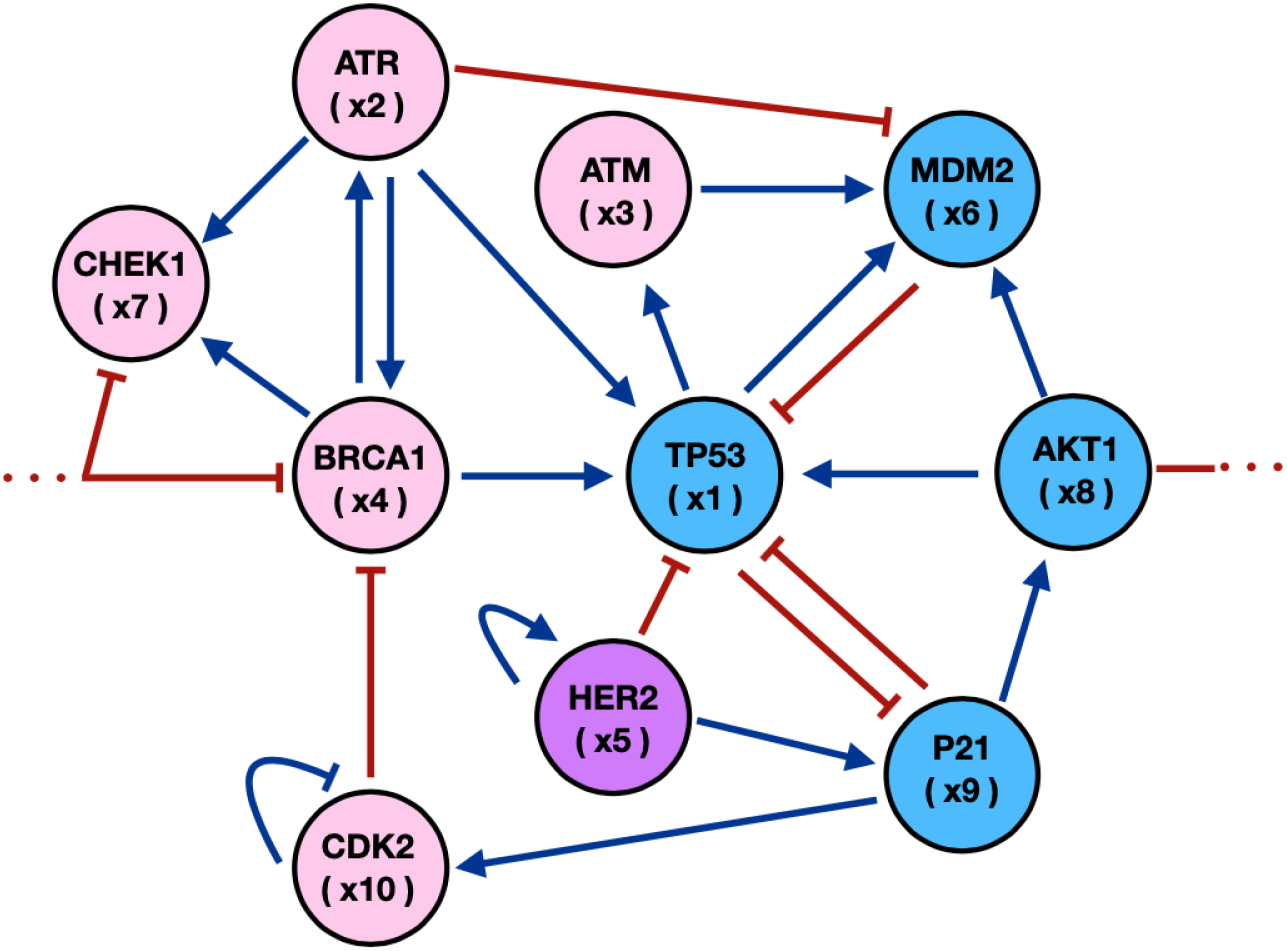
Gene Regulatory Network for Breast Cancer in which estrogens (pink) and oxygen (blue) concentrations in the microenvironment affect the gene states. The arrows indicate activation whereas the short bars represent inhibition interactions.

The kinetic interactions between genes are described by the following coupled system of nonlinear kinetic equations:

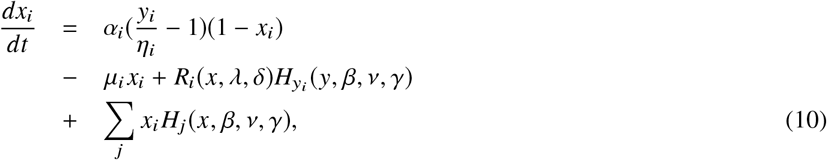

The function *x*_*i*_ represents the state of gene *i* (see Fig. 1), the parameters α_*i*_ represent the interaction rate constants between gene states and microenvironment. The action of the microenvironmental agents on gene activation is represented by *y*_*i*_ while η_*i*_ represents the activation-inhibition (A-I) threshold parameter and µ_*i*_ is the self-degradation constant. The parameters of Eqn. (10) were estimated by constructing and analyzing the Boolean representation of the GRN (see appendix A). In Table **??** of section 1.2 are indicated the specific values of these parameters and a brief explanation is given of how and why these values were chosen.

The self-regulatory positive (+) and negative (−) feedback genes interactions are given by the following equation,

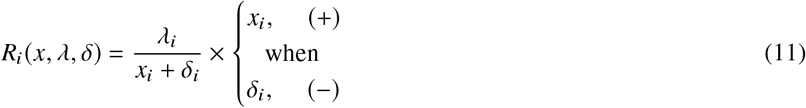

where λ_*i*_ is the activation constant and *δ*_*i*_ is the action threshold. Finally, the genetic interaction strength between genes is given by the Hill function.

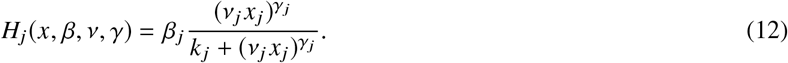

where *β*_*j*_ is the A-I strength, *ν*_*j*_ is the action velocity in which gene *j* affects the state of gene *i*. On the other hand, *k* _*j*_ represents the threshold of the sigmoidal function, *γ*_*j*_ is the Hill coefficient which depicts the steepness of the sigmoidal function representing the cooperativeness of the transcription factor regulatory binding to the genes. In the present case *k* _*j*_ = 1/ 2 for all genes, this means that all genes have the same chance to activate or inhibit each gene state. Moreover, the value of *γ*_*j*_ depends on how many neighbor genes change the state of gene *j*.

#### Microenvironmental action

We consider that oxygen and estrogens are the main microenvironmental factors that affect the GRN dynamics. In this context the variables *y*_1_, *y*_6_, *y*_8_ and *y*_9_ are determined by the normalized oxygen concentrations, while the remaining ones are proportional to the estrogen expression levels as,

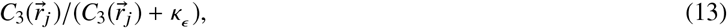

where *κ*_*ϵ*_ is the estrogens carrying capacity. The last equation emphasizes that GRN dynamics changes by the effective activation of estrogens levels. In this case, κ_*ϵ*_ regulates the threshold at which genes are expressed and wild type threshold values were chosen.

#### Intrinsic and extrinsic plasticity

To understand the intra-tumoral diversity that drives cancer development, the known facets of the disease can be categorized into cell-intrinsic and cell-extrinsic components. Intrinsic cell components are the inherent properties of a cell that contribute to its oncogenic phenotype as a result of collective molecular changes, stochastic genetic alterations as well as selective pressures. Extrinsic cell components are features related to micro-environmental variations that influence its phenotype perturbing the course of neoplastic disease (80). These intrinsic and extrinsic cell components can be categorized as cell intrinsic and extrinsic plasticity(81, 82).

A cell lineage is the developmental history of a differentiated cell as traced back to the cell from which it arose. Thus, cell linage depends on the ability to active or inhibit genes as a result of these two types of plasticity. As a first estimation of cell plasticity one can propose that gene activation-inhibition (A-I) is a linear superposition of the intrinsic and extrinsic plasticity. The former is proportional to the A-I parameter *η*_*i*_ while the latter is proportional to the segregation index 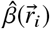. Thus, one can write down the following relationship for the variations of the threshold A-I parameter Δ*η*_*i*_.

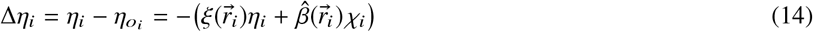

where η_*oi*_ is initial threshold value, 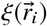 and χ are the intrinsic and the extrinsic (quorum sensing) susceptibilities, respectively. The minus sign in the right side of the equation indicates that there is certain opposition of the cell to change its plasticity. There are two limits in this equation, (i) when intrinsic plasticity (linear heritage) dominates genetic dynamics so that *ξ* becomes important, and quorum sensing susceptibility becomes negligible, and (ii) when extrinsic plasticity (epigenetic inheritance) dominates genetic dynamics and segregation becomes important whereas intrinsic plasticity is negligible.

### Numerical integration

The solutions of the reaction-diffusion system, Eqns (1) together with the death probability, Eqn. (2) and the cell cycle period for division, Eqn. (3), were calculated numerically. We assumed that normal, cancer and necrotic cells live on the sites of a square lattice of size *L*^2^ = 500^2^ (26) for 2D arrays and *L*^3^ = 225^3^ for 3D arrays. Nutrients, oxygen and estrogens were continuously supplied trough a capillary located at the top of the lattice simulating the bloodstream. Eqns. (1) were integrated using zero flow boundary conditions at the left, right, and lower sides of the 2D domain and on the four vertical and bottom sides of the cubic domain. In order to obtain an homogeneous diffusion of nutrients, oxygen and estrogens, the equations were solved locally for each node populated with cancer cells, using a grid of size 10^2^ units with zero flow boundary conditions, until the steady state was attained. After solving the equations locally the reaction-diffusion equations were solved globally until a simulation cycle was completed. A simulation cycle consists of a complete swap of the lattice, that is, once each site of the lattice has been visited.

At the beginning of the simulations all cells are normal except for one that has developed cancer and is located at around the lattice center. We assumed that this initial cancer cell has suffered mutations in the gene TP53, which is the gene that plays a crucial role in tumor’s growth. Once this initial cell begins to proliferate its descendants undergo mutations in all the other genes according to the probability distribution given in Eqn. (8). Once cell cycle is completed (using Eqn. 3), cell division occurs and the daughter cell position is chosen randomly with the same probability as either, one of the four nearest neighbors in the 2*D* array or one of the six nearest neighbors in the 3*D* array. If the site is occupied by a normal cell, it is replaced by the cancer cell, however, if the site is occupied by cancer cell, they stack. Then, a random number *r* distributed uniformly in the interval [0, 1] is chosen, and it is compared with the probability,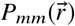, that a mutation occurs. See Eq (9). A mutation happens whenever,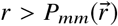, otherwise no mutation occurs and a new cycle is initiated by randomly choosing another cell and repeating the procedure described above for the mutations dynamics. We ran simulations to estimate the time in which the tumor reached the border of domain, since metastasis process was not considered in the model. For 2*D*, the largest tumor size was 450^2^, whereas a 3*D* the largest tumor size was 200^3^. Therefore, the required time to obtain the largest tumor was *T*_*max*_*≈*800 cycles. By using the duration of the cell cycle for breast cancer and normal cell (83–85) one can estimate, using Eq. (4), that each cell generation time *T* corresponds to 12-15 hours. We use a generation time to define one simulation cycle. To figure out the statistical meaning of the results we performed averages over 5, 10 and 20 simulations and found that the results were consistent with those corresponding to one simulation within one standard deviation. Therefore, most of the results reported here correspond to one simulation of the system. The values of the simulations parameters were chosen in accordance with reference (26). That is, we considered *θ*_div_ = 0.3, *θ*_dead_ = 0.01, λ_1_ = {50, 100, 200}, λ_2_ = 50, λ_3_ = {50, 100, 200}, α_1_ = {2/ *L*, 4 / *L*, 8/ *L*} and α_3_ ={2/ *L*, 4/ *L*, 8/ *L*}. By recalling that in the present model the probability *P* (*B*) is considered as a varying parameter. Thus, we performed simulations for *P* (*B*) = 0.05, 0.1, 0.25, 0.5, 0.75, and 1.0. The results and their corresponding interpretation and discussion are presented in the following section.

## RESULTS

Microarrays may be used to measure gene expression in many ways, but one of the most popular applications is to compare expression of a set of genes from a cell maintained in a particular condition to the same set of genes from a reference cell maintained under normal conditions (wild type). Bearing this in mind, here we present our findings for genes expressions in terms of series of microarrays that facilitate the characterization of the different states of the GRN shown in Fig. 1. These states were obtained from the numerical solutions of the equations that describe the GRN dynamics. See Eqns. (10-13). The microarrays presented in Figs. 2 and 3 show two vertical axes. On the left vertical axis is indicated each one of the ten genes that are part of the GRN, while on the right vertical axis is indicated the micro-environmental agent, either oxygen or estrogens, that contributes to the expression of the corresponding gene. The horizontal axis indicates the normalized concentration of each microenvironmental agent.

**Figure 2:**
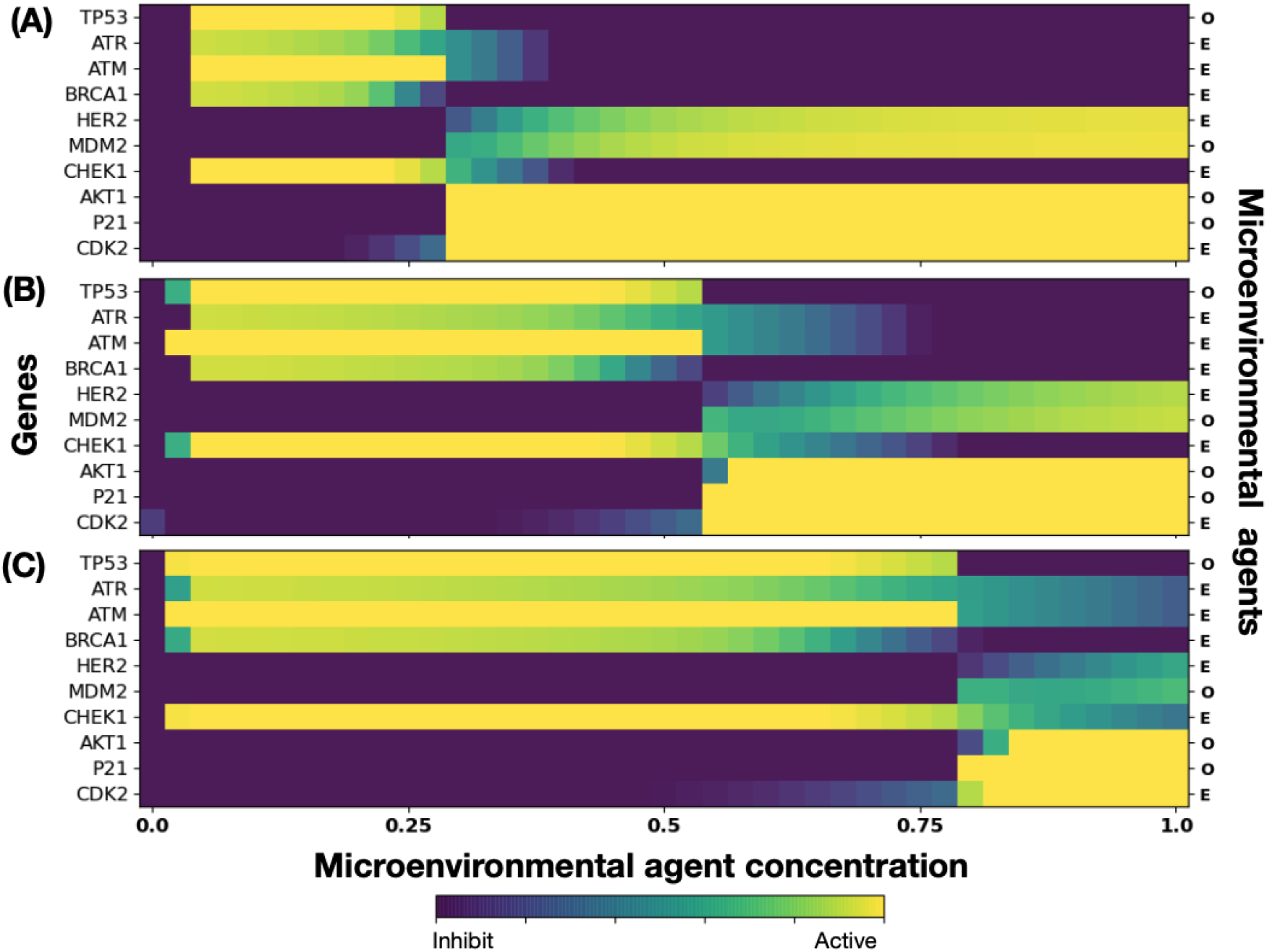
Microarray that describes the genes expression levels of the GRN in terms of the concentrations of glucose, oxygen (O) and estrogens (E) for different initial values of the activation/inhibition (A-I) threshold parameters. (A) 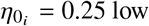, (B) 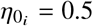medium, and (C) 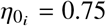 moderate.

**Figure 3:**
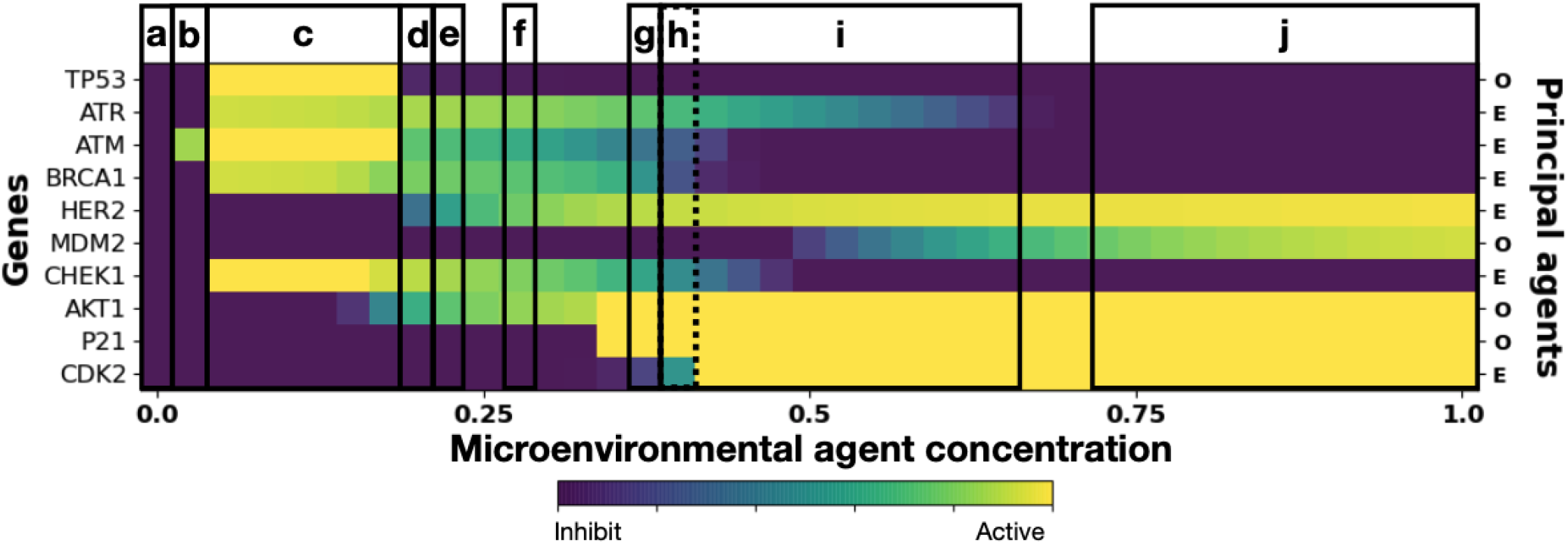
Microarrays that shows genes expression and cell phenotypes. Normal cell phenotypes are represented by the states **a** and **c**. The states **b, d, e, f, g, h**, and **i** are identified with attractors in which some genes are Over-Expressed or Knocked-Out, suggesting a precancer phenotype. The state **j** is identified with the attractor that represents cancer phenotypes. The initial threshold parameters for the set of ten genes are: 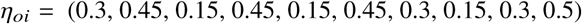. The phenotypes are associated with the GRN dynamics –see Fig. 1– that can also be described by a continuous model (see appendix B).

As an initial test to the response of the GRN to different initial values of the A-I threshold parameter 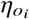, in Fig. 2 are presented three microarrays that correspond to the following three different values: A) 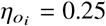, low activation, B) 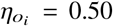, medium activation, and C) 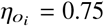, moderate activation, for all genes, respectively. We begin by observing that, as expected, all genes in the GRN need a nonzero normalized concentration value for at least one of the genes be expressed by one of the environmental agents. This concentration value depends on the gene in turn, for instance, in Fig. 2(A), genes HER2, MDM2, AKT1, P21 and CDK2 need a concentration higher than 30% to be expressed. These concentrations increase in about the same proportion as the initial A-I threshold parameter is increased. Observe that genes TP53, ATR, ATM and BRCA1 are expressed at relatively low concentrations in a well defined concentration window whose size increases as the initial A-I threshold parameter increases. Note that for 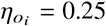 genes AKT1, P21 and CDK2 are fully expressed for oxygen and estrogens concentrations greater than 0.25, whereas for 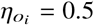 and 0.75 these genes are fully expressed for concentrations greater than 0.50. Nonetheless, genes ATR, ATM, BRCA1, MDM2 and CHEK1, are expressed for oxygen and estrogens concentrations smaller than 0.50. The expression of each gene is shown in the microarray of Fig. 2(B) when the initial values of 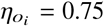 for all genes is set at the middle saturation value 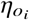. There one sees that genes TP53, ATR, ATM, BRCA1, CHEK1 and CDK2 become active for oxygen and estrogens concentrations less than 0.5, However, genes HER2 and MDM2 become active for estrogen and oxygen concentrations greater than 0.75, respectively. Similarly, genes AKT1, P21 and CDK2 become activated for oxygen and estrogen concentrations greater than 0.50. These results indicate that the first four genes dominate the GRN dynamics for 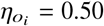. In the microarray of Fig. 2(C) obtained for 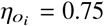 for all genes, one sees that genes TP53, ATR, ATM, BRCA1, CHEK1 and CDK2 become activated for oxygen and estrogens concentrations less than 0.75. Nonetheless, genes HER2, MDM2, ATK1,P21 and CDK2 become activated for estrogen and oxygen concentrations greater than 0.80, respectively. These results indicate that for moderate values of 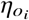 genes TP53, ATR, ATM, BRCA1 and CHEK1, dominate, once more, the GRN dynamics.

Figure 3 shows microarrays associated to three cell phenotypes during cancer development: (i) normal cells, (ii) precancerous cells, and (iii) cancer cells. These phenotypes were identified as attractors in the dynamics described by the GRN shown in Fig. 1. GRN dynamics was analyzed by both, a continuous model, Eqns. (10), and a Boolean representation of the genes (see appendix B) yielding consistent results. Attractor states **a** and **c** are identified with normal phenotypes because in **a** all genes are inhibited while in **c** only genes TP53, ATM and CHECK1 are over-expressed. However, attractor states **b, d, e, f, g, h**, and **i** can be identified in the Boolean GRN representation with a precancer phenotype in which some genes are either over-expressed or inhibited permanently. Finally, the state **j** is the only attractor that one identifies with cancer phenotype because genes HER2, ATK1, P21, and CDK2 are over-expressed. These results are in agreement with a previous GRN analysis (86). The other hand, we estimated the effect of estrogens on gene expression levels by varying the activation thresholds, 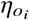, for each gene proportionally and by making a variation of only the gene HER2 (see Appendix B). The results show that the precancer phenotype decreases more when an increase in estrogens initial threshold values occurs in gene HER2 rather than in all other genes. This results is in agreement with the acceleration of tumorigenesis reported in Ref. (24) when HER2 is overexpressed and also reaffirms the role of estrogen concentration in changing the gene expression levels and the occurrence of different phenotypes as previous findings have shown (17, 18, 20, 56, 70).

Figure 4 shows the spatial distribution of cell phenotypes -panel (A)– and segregation index –panel (B)– obtained from simulations of a 2D tumor that incorporate the mutation dynamics described in section, together with the GRN dynamics with the A-I threshold values of all genes set at the middle saturation values 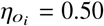. The values of the kinetic parameters were carefully chosen so that the three attractor states associated with the cell phenotypes were obtained. In Fig. 4(A) we observe tumor regions populated with: (i) cells in a normal state (purple regions), (ii) cells in a precancer state green regions, and (iii) cells in a cancer state (yellow regions). Fig. 4(B) shows the genetic cell lineages spatial distribution identified through their segregation index values. On the right side of the figure is shown –vertical bar– the color scale of the segregation index. One should recall that the segregation index directly measures the spatial assortment of genetic cell lineages and ranges from 0 to 1, where 1 denotes complete lineage segregation on a given spatial scale. The segregation index 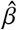 was calculated for 2D and 3D tumors using Eqn. 5 on grids of size 20^2^ and 20^3^, respectively. Notice that the tumor upper region is populated with cancer cells and it is precisely in this region where the segregation index values are greater than 0.8 as an indication of cancer cells segregation and genetic heterogeneity. On the contrary, the tumor lower region is mostly populated by normal cells with segregation index values relatively low, 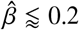. At the middle of the tumor, in a smaller region –as compared to the other two phenotypes–, precancer cells phenotype is located and the corresponding segregation index values, 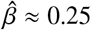 can be considered as moderate.

**Figure 4:**
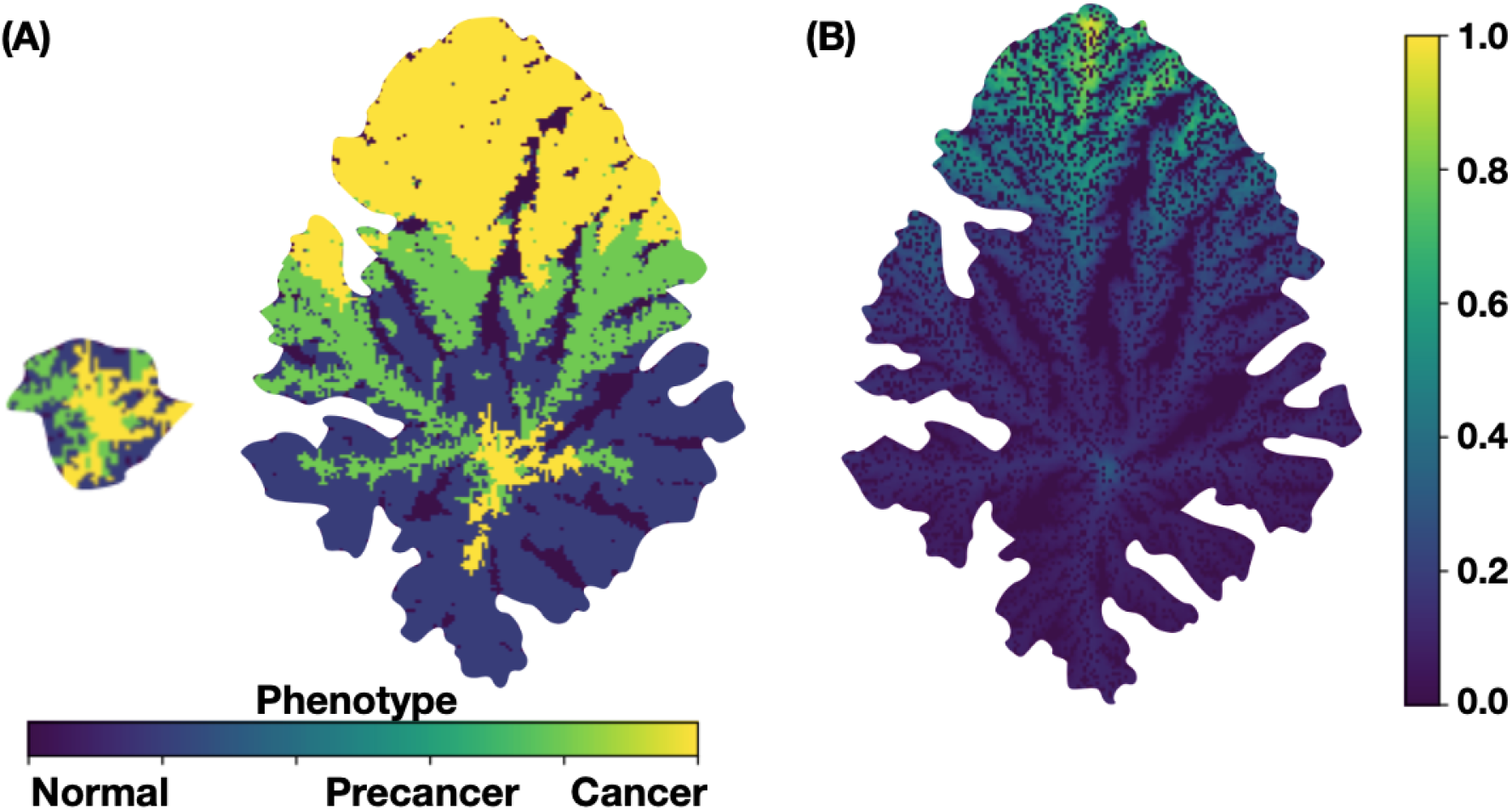
For a 2D tumor the spatial distribution of cell phenotypes (A) and Segregation index (B) are shown. These results were obtained for the model parameters values : *P* (*B*) = 1, κ_*ϵ*_ = 0.1, *ξ* = 0.5, χ = 4, α_1_ = α_2_ = *α*_3_ = 8× 10^−3^, *λ*_1_ = 100, *λ*_2_ = 50, and *λ*_3_ = 200.

Figure 5 shows the spatial distribution of mutations for a 2D –panel (A)– and for a 3D –panel (B)– tumors at two times of evolution stage. The upper figures of both panels represent tumors at about two months of development while the lower figures correspond to the same tumors after one year of evolution. Notice that both tumors developed a fractal-like structure at the early stages whereas they developed a solid-like structure at the later stages. The colorbar indicates the spatial distribution of the number of mutations in the cancer cells. One should also observe that genetic heterogeneity is more marked at the early stage of development as compared to the later stage. The regions where number of mutations is maximum are relatively small and they occur at the tumor surface. These results were obtained using the same kinetic parameters values as in Figure 4.

**Figure 5:**
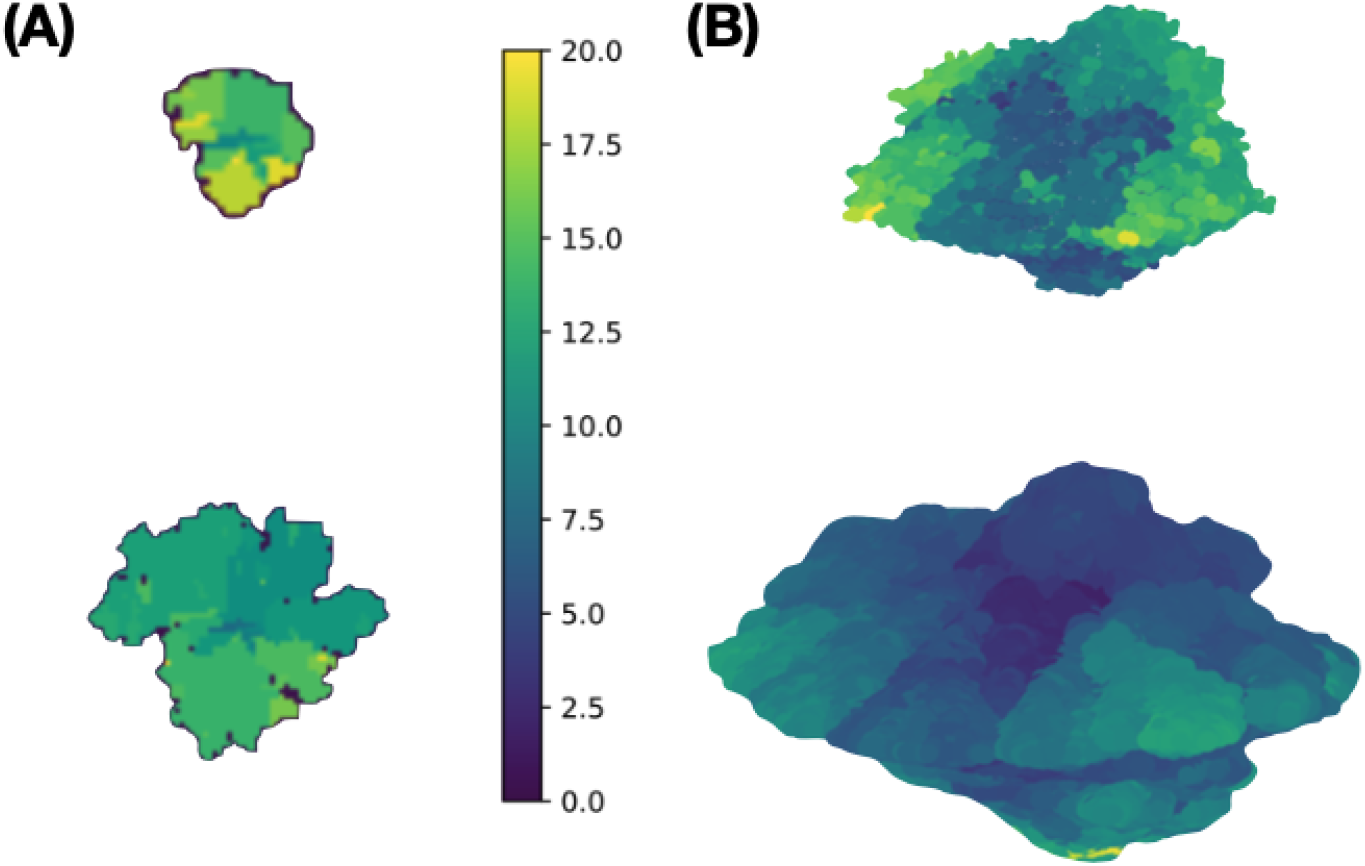
Spatial distribution of the number of mutations for (A) 2D and (B) 3D tumors. These results were obtained for the same model parameters values as in Fig. 4.

Figure 6 shows four microarrays that illustrate the estrogens carrying capacity, κ_*ϵ*_, proposed for the continuous dynamics of the GRN and described by Eq.(14).

**Figure 6:**
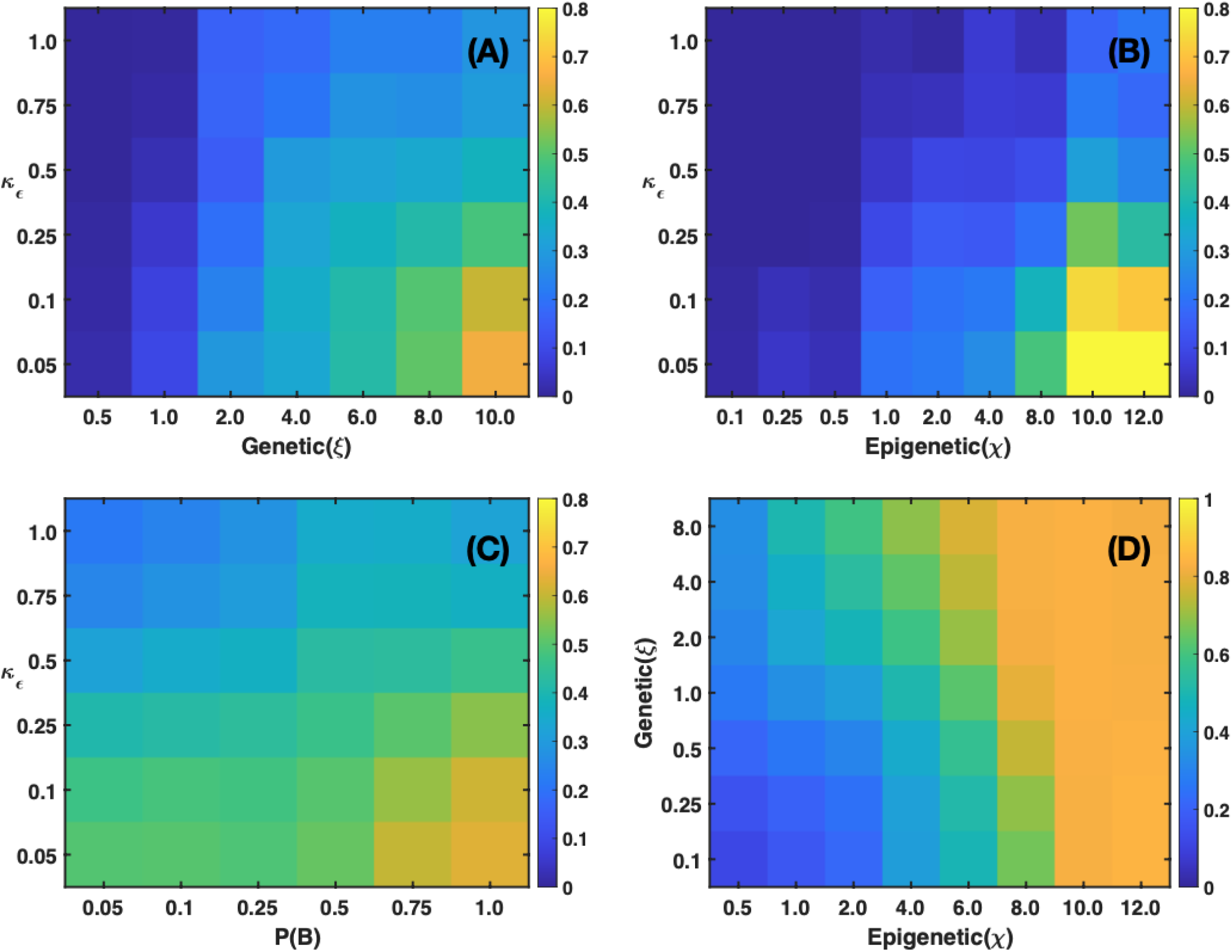
The microarrays show the fraction of cancer cells in a 2D tumor for (A) Estrogens expression levels versus genetic inheritance values. (B) Estrogens expression levels versus epigenetic inheritance values. (C) Estrogens expression levels versus estrogen receptors probability. (D) Genetic inheritance versus epigenetic inheritance. The microarrays were obtained with the following parameter values: (A) *P*(*B*) = 1 and χ = 0; (B) *P*(*B*) = 1 and *ξ* = 0; (C) χ = 4 and *ξ* = 0.5; (D) *P*(*B*) = 1 and κ_*ϵ*_ = 0.1. The consumption parameter values were *α*_1_ = *α*_2_ = *α*_3_ = 8 × 10^−3^, *λ*_1_ = 100, *λ*_2_ = 50, and *λ*_3_ = 200.

Figure 6(A) shows a microarray obtained for a 2D tumor that was built for different values of the estrogens cooperative activation, κ_*ϵ*_ (vertical axis), as well as different values of genetic heritage, *ξ* (horizontal axis). It is seen that for certain ranges of κ_*ϵ*_ and *ξ* there occurs a full estrogens expression that leads to an increasing number of cancer cells. Nonetheless, for the complementary ranges of these parameters there is a full estrogen inhibition. In figure 6(B), the microarray indicates that epigenetic inheritance, χ, follows a similar genetic heritage behavior as a function of κ_*ϵ*_. These results demonstrate that genetic and epigenetic inheritance may change the emergence of different cancer phenotypes. Because of this, one would expect that there exist a mechanism that can revert the epigenetic changes. The microarray presented in figure 6(C), shows the importance of the probability *P*(*B*) due to failure of estrogens receptors for different values of κ_*ϵ*_. This microarray suggests that if cells developed the ability to keep activated a high number of estrogen receptors then the emergence of cancer phenotypes would be reduced. The microarray presented in figure 6(D) shows the genetic versus epigenetic heritages. The results suggest that the occurrence of cancer phenotypes in the tumor is the result of epigenetic changes rather than genetic changes. Because of this one may think of the possibility of decreasing epigenetic changes, and therefore, the number of cancer phenotypes by manipulating the concentrations of microenvironmental substances –nutrients, estrogens and drugs–.

In Fig. 7, the microarrays explore the importance of genetic and epigenetic heritages in the development of 2D tumors. The microarray in figure 7(A) indicates that the increase in segregation index correlates with epigenetic changes. This suggests that genetic expression thresholds are more susceptible to changes in quorum sensing rather than to random mutations derived from nutrients and estrogens concentration gradients. Figure 7(B) shows that the fraction of normal cells in a tumor is relatively high when epigenetic and genetic inheritances are small, suggesting that these quantities play similar roles. Microarrays in figures 7(C) and 7(D) show the variations in the accumulation of mutations and heterogeneity indexes for different values of the genetic and epigenetic heritages. It can also be observed that epigenetic heritage is more relevant as in figure 7(A). The present results strongly suggest that variations in epigenetic inheritance define the degree of tumor malignancy, in agreement with previous results (26). These findings also suggest that it should be possible to reverse cancer phenotypes by manipulating the microenvironmental agent concentrations, as suggested by several authors (17, 18, 20, 56, 70).

**Figure 7:**
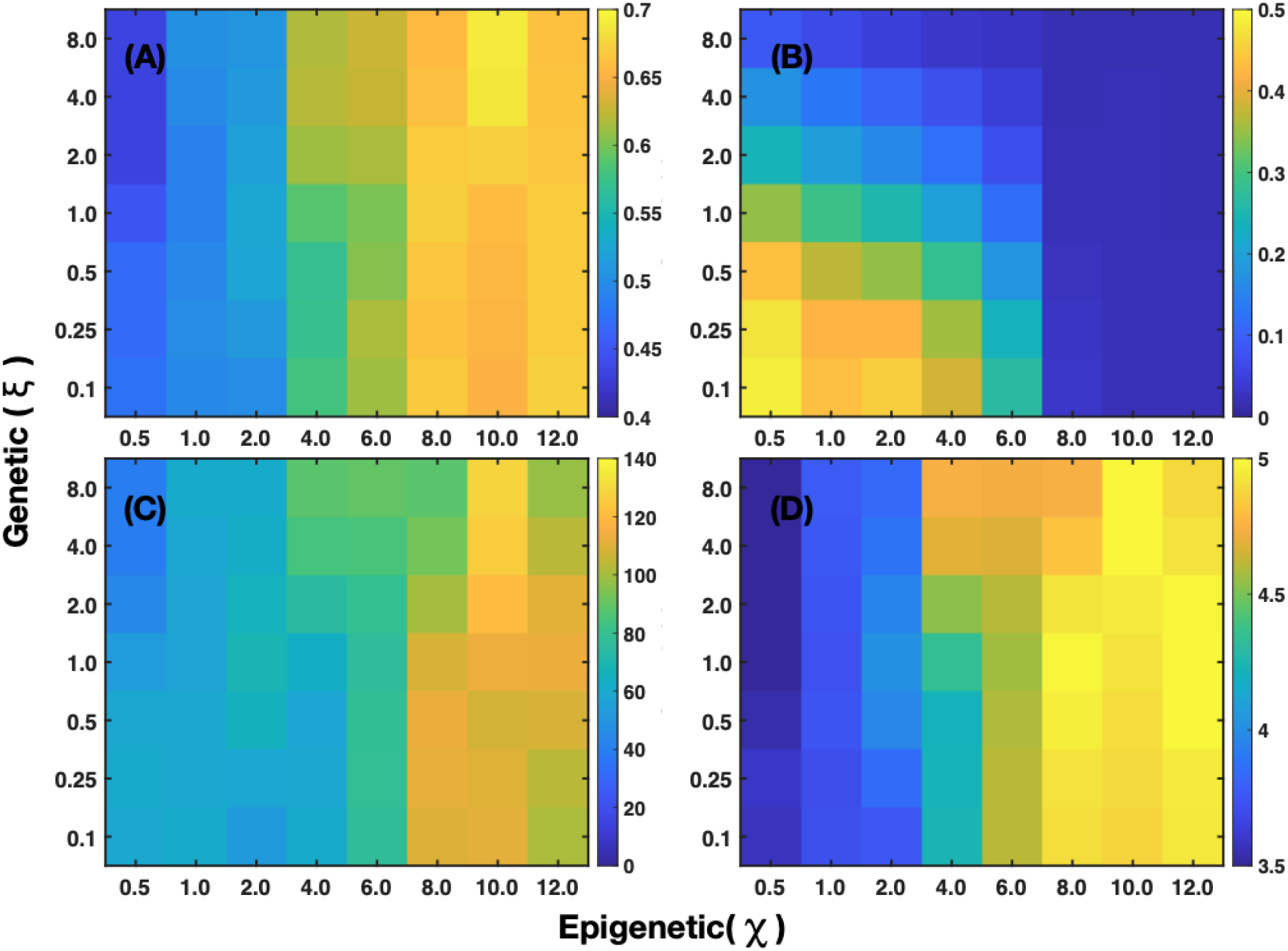
The microarrays correspond to a 2D tumor and represent: (A) Mean segregation index. (B) Fraction of normal cells in a tumor, (C) Average number of mutations. (D) Shannon index that measures tumor heterogeneity. These microarrays were obtained for the following parameter values: *P*(*B*) = 1, κ_*ϵ*_ = 0.1, *α*_1_ = *α*_2_ = *α*_3_ = 8 × 10^−3^, *λ*_1_ = 100, *λ*_2_ = 50 and *λ*_3_ = 200.

Figure 8 shows the fraction of cancer cells, the segregation index spatial distribution, different tumor shapes, as well as the fractal and Shannon indices obtained for a 2D tumor. Microarrays (A), (C) and (D) consist of a main lattice (heavy lines) that indicates the estrogens consumption rate, *α*_3_ (vertical axis) and the estrogens supply rate, *λ*_3_ (horizontal axis). Inside each cell of the main lattice there is a sublattice that indicates the distribution in terms of nutrients consumption rate, *α*_1_ (vertical axis) and the nutrients supply rate, *λ*_1_ (horizontal axis). Panel (A) shows how the fraction of cancer cells is distributed throughout the tumor. This microarray suggests that cancer phenotype is mostly driven by estrogens consumption rather than nutrients consumption. Nonetheless, precancer phenotype happens for low (lower left square) and intermediate (centered square) values of estrogens consumption and supply rates. Normal phenotype is present for relatively high (upper left square) and intermediate (upper centered square) values of estrogens consumption and supply rates. Panel (B) shows the spatial distribution of segregation index throughout the tumor as well as different tumor shapes that are different depending on the values of the estrogens and nutrients consumption and supply rates. It is observed that segregation maximizes in the regions where estrogens gradient concentrations is large. In addition, it can be observed that at the tumor center and contour genetic diversity increases too. The parameters values to obtain the tumors in panel (B) correspond to the center values of the microarrays in panel (A). Note that the fractal tumor shapes shown in the third column top and middle rows correspond to *α*_1_ = 8 ×10^−3^, and *λ*_1_ = 100 and *α*_3_ = 8× 10^−3^, 16 ×10^−3^ with *λ*_3_ = 200. Panel (C) shows the fractal index distribution where it is observed that its higher values occur for *α*_1_ ∼ 4 ×10^−3^ and for 100 ≤ *λ*_3_ ≤ 200, pointing to a malignant or cancer cell phenotype, while the lower values happen for *α*_1_∼ 16× 10^−3^ and for values of *λ*_3_∼ 50 suggesting the presence of a precancer cell phenotype. These results are consistent with those shown in panel (A) and those reported in reference (26). The Shannon index distribution is shown in panel (D). The higher values of this index occur for *α*_1_ ∼4 ×10^−3^ and for 100 ≤ *λ*_3_ ≤ 200, pointing to a malignant or cancer cell phenotype. The results shown in this panel suggest that during cancer evolution estrogens consumption is more relevant than nutrients consumption which means that the increase in tumor diversity is mainly due to the presence of estrogens concentration gradients. The results in the sublattice are also consistent with those shown in panels (A)-(C) and those reported in reference (26) for nutrients consumption.

**Figure 8:**
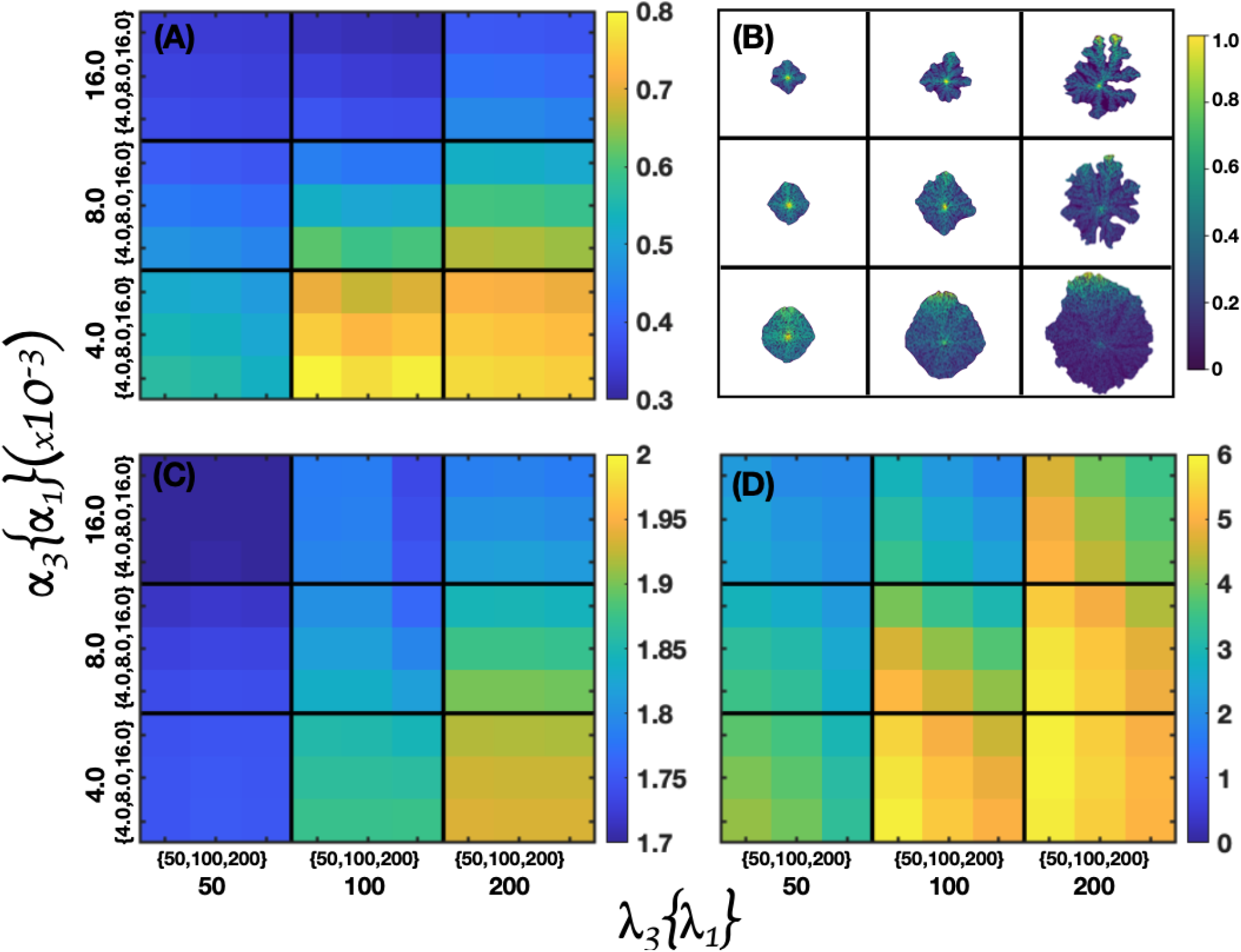
Microarrays that correspond to a 2D tumor. (A) Fraction of cancer cells. (B) Tumor shape and spatial distribution of segregation index. (C) Fractal index, D) Shannon index. The microarrays were obtained for the following parameter values: *P*(*B*) = 1, κ_*ϵ*_ = 0.1, *ξ* = 0.5 and χ = 4.

Figure 9 presents four panels corresponding to three microarrays that explore the importance of genetic and epigenetic heritages, and the spatial distribution of random mutations in the development of 3D tumors. The microarray in figure 9(A) shows that the increase in the fraction of cancer cells is correlated with the epigenetic changes suggesting that genetic expression is more susceptible to quorum sensing than to random mutations. Figure 9(B) shows the spatial distribution of random mutations in 3D tumors under different microenvironmental conditions defined by the estrogens consumption and supply rate values. There one sees that for *α*_3_ = 16 × 10^−3^ and *λ*_3_ = 50, and 100 there are relatively small regions in the tumor where the the number of mutations is of the order of one thousand. However, for *α*_3_ = 8× 10^−3^ and 16× 10^−3^ and *λ*_3_ = 200, there are large regions in the tumor where the the number of mutations is of the order of one hundred. For *α*_3_ = 4× 10^−3^ and *λ*_3_ = 100 and 200, there are relatively large regions in the tumor that undergo between one thousand and fifteen hundred mutations. It was also found that cells that underwent more mutations were located close to the tumor surface. This result is consistent with that found for the 2D model in which most mutations were found at the perimeter of the tumor. One can also observe that the interior regions of the 2D and 3D tumors were mostly populated with necrotic cells. The microarray in figure 9(C) shows the effect of varying the genetic and epigenetic heritage threshold values whereas the panel in figure 9(D) shows the distribution of the Shannon index values as genetic and epigenetic threshold values are varied. As seen in panel 9(A), epigenetic effects are more relevant, which suggests that a decrease in the epigenetic threshold values leads to an increase in tumor heterogeneity. By examining the features presented in Fig. 7 for a 2D tumor and in Fig. 9 for an equivalent 3D tumor it can concluded that all their characteristics are consistent with each other.

**Figure 9:**
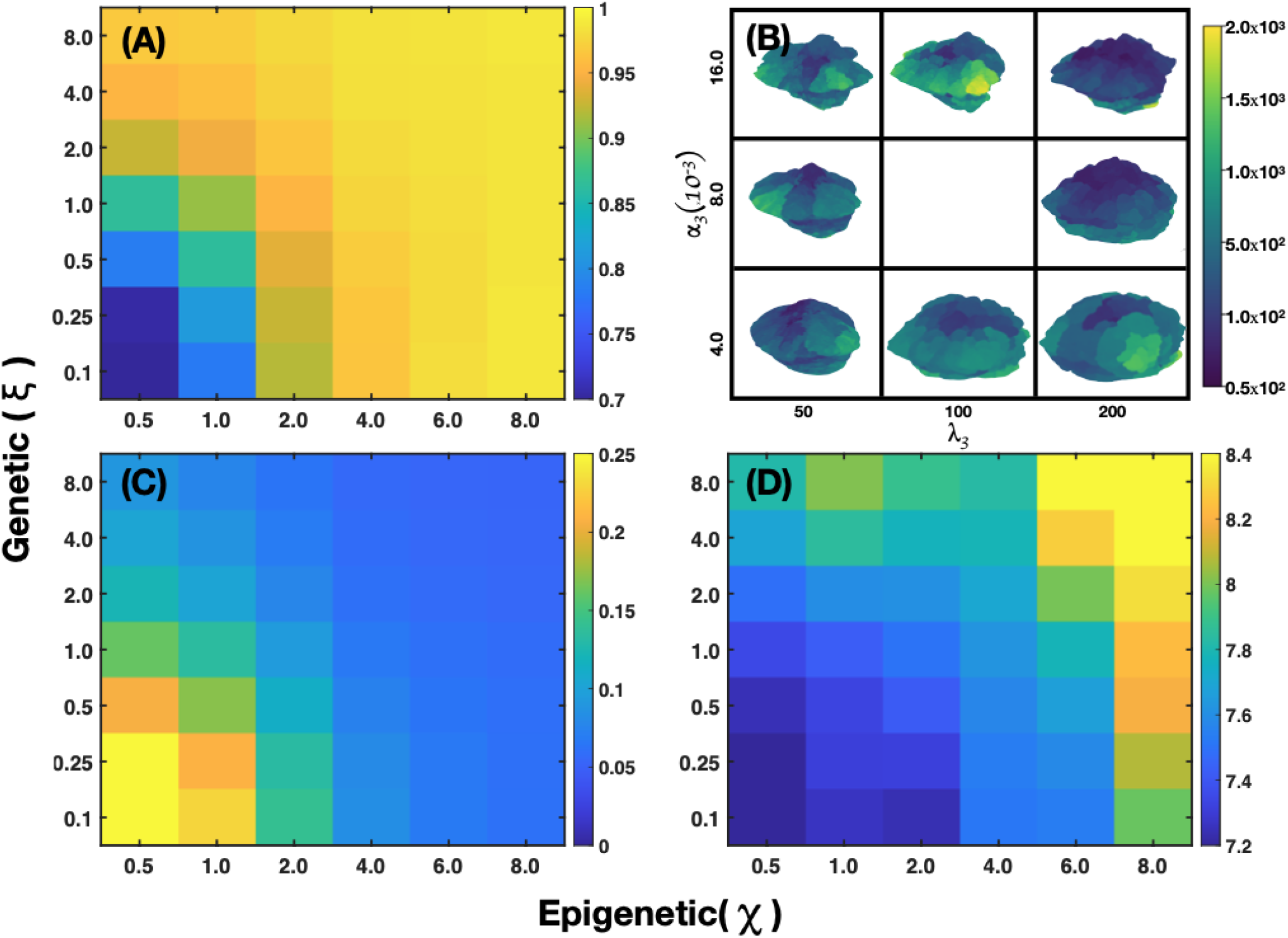
Microarrays obtained for 3D tumors that show: (A) Fraction of cancer cells, (B) The spatial distribution of mutations for different values of *α*_3_ and *λ*_3_. (C) Average threshold value, (D) Tumor heterogeneity. The results shown in (A), (C) and (D) were obtained for the following parameter values: *P* (*B*) = 1, κ_*ϵ*_ = 0.1, *α*_1_ = *α*_2_ = *α*_3_ = 8 ×10^−3^, *λ*_1_ = 100, *λ*_2_ = 50, and *λ*_3_ = 200.

Figure 10 presents the distribution of phenotypes, for a 2D tumor in panel (A) and for a 3D tumor in panel (B), for different values of the estrogen consumption *α*_3_ and supply rates *λ*_3_, while maintaining all the other model parameter values as in Fig. 9. Panel (A) shows that in the column corresponding to *λ*_3_= 50 the tumor looks compact and its size increases as *α*_3_ decreases. For intermediate and higher values of this quantity precancer and cancer phenotypes can be observed. As *λ*_3_ increases the tumor develops a fractal like structure and the three phenotypes can be clearly observed for the highest values of the estrogen consumption and supply rates. Panel (B) shows 3D tumor structures and phenotype spatial distributions for different values of the estrogen consumption *α*_3_ and supply rates *λ*_3_ which are consistent with those observed in panel (A). In the third column that correspond to the highest value of *λ*_3_, and the three values of *α*_3_, the distributions of normal, precancer and cancer phenotypes as well as the tumor fractal like structure are more evident. The tumor surface is mostly populated by cancer and precancer phenotypes whereas small regions are populated by normal phenotype.

**Figure 10:**
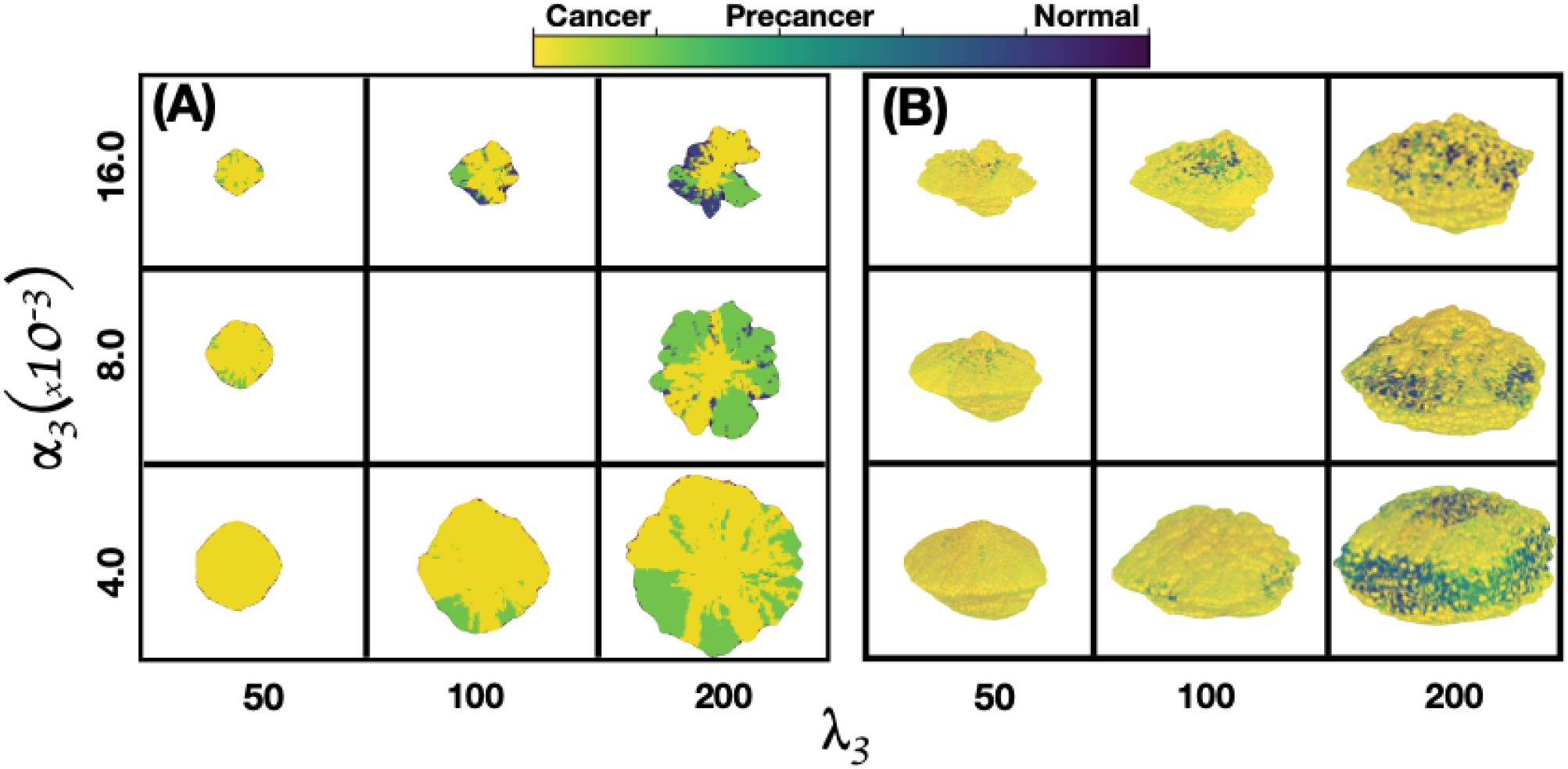
Spatial distribution of phenotypes for different values of the estrogen consumption *α*_3_ and supply rates *λ*_3_. (A) 2D tumor and (B) 3D tumor. All the model parameters are the same as in Fig. 9.

## CONCLUSIONS

We have introduced and analyzed in detail a quantitative model that describes the growth and epigenetic evolution of an avascular tumor. The epigenetics was described through the dynamics of a GRN formed by the set of ten genes: TP53, ATM, HER2, BRCA1, AKT1, ATR, CHEK1, MDM2, CDK2, P21 that are believed to play an important role in breast cancer development. The GRN dynamics was analyzed using a Boolean representation of the genes and their interactions as well as by means of a continuous model described by ten kinetic equations that involve transcriptional positive and negative regulations. The tumor growth was simulated with a cellular automata coupled to a set of reaction-diffusion equations that described the transport of the microenvironmental agents glucose, oxygen and estrogens that changed the genes expression levels. Random mutations were simulated by means of a Markovian process modeled with a master equation that involved the local concentrations of the microenvironmental agents. The role of cell segregation was also incorporated by modeling the lineage production rate –cell segregation index– with a metabolic Michaeles-Menten kinetic equation. The lineage production rate was introduced as a factor in the probability of mutations associated to the genotoxic metabolites driven by estrogen concentration gradients. This approach led us to find three attractors in the GRN dynamics which are related to three phenotypes: (i) normal cells, (ii) precancer cells and (iii) cancer cells. With these ingredients the tumor structure, the spatial distribution of mutations and phenotypes for 2D and 3D tumors were calculated. From the simulations we obtained a series of microarrays that describe the activation levels of each of the ten genes in response to the nutrients and estrogen concentration gradients. In addition, we obtained the spatial distributions of: (i) the number of mutations, (ii) the mutations, (iii) phenotypes, (iv) shape and, (v) the Shannon index, that is a measure of the heterogeneity for 2D and 3D tumors. These quantities were also analyzed for different values of the estrogen consumption and supply rates. It was found that the regions where number of mutations maximizes are relatively small and they occur at the tumor surface. whereas genetic heterogeneity was more marked at the early stage of development. The segregation index spatial distribution throughout the tumor as well as different tumor shapes were different depending on the values of the estrogens and nutrients consumption and supply rates. It was also found that segregation maximizes in the regions where estrogens gradient concentrations are large.

Tumors developed a fractal-like structure at the early stages whereas at the later stages they tended to develop a solid-like structure. On the other hand, we studied the role of estrogen concentration in changing the gene expression levels and the results reaffirm that the phenotypes can be controled directly by estrogen concentration. All these results were found to be consistent for both, 2D and 3D tumors. The findings reported here strongly suggest that it is possible to develop epigenetic cancer treatment alternatives. Finally, we would like to emphasize that the results obtained for the fractal structure and heterogeneity of tumors are in complete agreement with those found in a previous paper (26)

## ACKNOWLEDGMENTS

JRRA thanks the IIMAS Department of Mathematics and Mechanics for allowing him to join its group of researchers. Likewise, JRRA thanks to Ana Pérez Arteaga and Ramiro Chávez Tovar for their technical support. GRS received support from DAGAPA-UNAM grant No. IN-109722. We thank the IIMAS and Laboratorio Universitario de Cómputo de Alto Rendimiento (LUCAR) for the cluster allocated CPU time.

## SUPPLEMENTARY MATERIAL

### APPENDIX A. GENE FUNCTIONS

This section describes the functional actions (obtained from Ref. (87)) of the ten genes that are part of the GRN shown in Fig. 1. This set of genes are believed to play an important role in breast cancer evolution.

**TP53(***X*_1_): Tumor suppressor gene, It controls the cellular genome’s integrity and is an important regulator of cell cycling, proliferation, apoptosis and metabolism. Mutations of TP53 or inactivation of its genetic product are among the first events that give rise to malignant cell transformation. The loss of control over the cell cycle, results in the acceleration of cell proliferation and facilitates metabolic reprogramming, giving the pre-malignant cells numerous advantages over healthy cells. Protein p53 becomes activated in response to countless stressors, for instance, DNA damage (induced by either UV, IR, or chemical agents such as hydrogen peroxide), oxidative stress, osmotic shock, ribonucleotide depletion, and deregulated oncogene expression. The activation of protein p53 is characterized by two major events. (i) A drastic increase of the half-life of the protein p53 leads to a quick accumulation of p53 in stressed cells. (ii) A conformational change in protein p53 drives its activation as a transcription regulator in the cell.

**ATR(***X*_2_): Kinase, It functions to maintain genome integrity by stabilizing replication forks and by regulating cell cycle progression and DNA repair. ATR is activated in response to persistent single-stranded DNA, which is a common intermediate formed during DNA damage detection and repair.

**ATM(***X*_3_): Kinase, It is a protein that coordinates DNA repair by activating enzymes that fix the broken strands. Efficient repair of damaged DNA strands helps maintain the stability of the cell’s genetic information. A wealth of evidence has accumulated showing that ATM is part of many signaling networks that include cell metabolism and growth, oxidative stress, and chromatin remodeling, all of which favor cancer progression.

**BRCA1(***X*_4_): Oncogene, It controls cellular pathways that maintain the genome stability, including DNA damage-induced cell cycle checkpoint activation, DNA damage repair, as well as transcriptional regulation and apoptosis.

**HER2(***X*_5_): Oncogene, when HER2 is normally expressed, ligands that bind to the HER receptors form only a few HER2 heterodimers, and the responses to these growth factors are relatively weak, resulting in normal growth of cells. However, when HER2 is over expressed as in cancer cells, many ligands originating primarily in the stroma or in the tumor cells themselves will recruit HER2 into heterodimers. The heterodimers stay longer at the cell surface, and their signaling is enhanced. This results in potent stroma-to-epithelium signaling, enhanced responsiveness to growth factors and, eventually, malignant growth.

**MDM2(***X*_6_): Oncogene, It is a key negative regulator of p53 protein and forms an auto-regulatory feedback loop with p53. MDM2 is highly regulated; the levels and function of MDM2 are regulated at the transcriptional, translational and post-translational levels. Following DNA damage, phosphorylation of MDM2 leads to changes in protein function and stabilization of p53.

**CHEK1(***X*_7_): Kinase, It is a protein kinase that has a central role in coordinating DNA damage response and cell cycle checkpoint response. Activation of CHEK1 results in the initiation of cell cycle checkpoints, cell cycle arrest, DNA repair and cell death to prevent damaged cells from progressing through the cell cycle.

**AKT1(***X*_8_): Kinase, It is one of three closely related serine/threonine-protein kinases (AKT1, AKT2, and AKT3) that regulates many processes including metabolism, proliferation, cell survival, growth, and angiogenesis by phosphorylating a range of downstream substrates in response to growth factor stimulation.

**P21(***X*_9_): Tumor suppressor gene, It is a cyclin-dependent kinase inhibitor of the tumour suppressor p53. It mediates cell cycle arrest in the G1 phase and cell senescence in response to several stimuli, including oncogene-induced proliferation. However, p21 can be activated independently of p53 and has cancer-promoting properties, such as supporting tumour growth. P21 is responsible for a bifurcation in CDK2 activity following mitosis, cells with high P21 enter a G0/quiescent state, whilst those with low P21 continue to proliferate. P21 may inhibit apoptosis in response to replication fork stress, however, it does not induce cell death on its own.

**CDK2(***X*_10_): Kinase, This gene encodes a member of a family of serine/threonine protein kinases that participate in cell cycle regulation. The encoded protein is the catalytic subunit of the cyclin-dependent protein kinase complex, which regulates progression through the cell cycle. Activity of this protein is especially critical during the G1 to S phase transition.

### 1 APPENDIX B. EPIGENETIC ATTRACTORS. GRN BOOLEAN AND CONTINUOUS DYNAMICS

#### 1.1 Boolean model

To identify how different genes interact with each other in the breast cancer GRN, in the context of microenvironmental agents, a Boolean network model was developed and analyzed. The interaction rules were constructed on the basis of the biophysical interactions reported by Chong Yu and Jin Wang (86) and are represented by the links depicted in Figure 1. The Boolean functions used to analyze the network model are given in logical Table (1).

**Table 1:** Network Boolean rules for breast cancer development.

| Gene | Boolean rules |
| --- | --- |
| **TP53**($x_1$) | $x_2 \wedge x_4 \wedge \neg x_5 \wedge \neg x_6 \wedge x_8 \neg x_9$ |
| **ATR**($x_2$) | $x_4$ |
| **ATM**($x_3$) | $x_1$ |
| **BRAC1**($x_4$) | $x_2 \wedge \neg x_8 \wedge \neg x_{10}$ |
| **HER2**($x_5$) | $x_5$ |
| **MDM2**($x_6$) | $x_1 \wedge \neg x_2 \wedge x_3 \wedge x_8$ |
| **CHEK1**($x_7$) | $x_9$ |
| **AKT1**($x_8$) | $x_2 \wedge x_4 \wedge \neg x_8$ |
| **P21**($x_9$) | $\neg x_1 \wedge x_5$ |
| **CDK2**($x_{10}$) | $x_9 \wedge \neg x_{10}$ |

A Boolean network model is the simplest formalism for studying the nonlinear interactions among genes and agents that constitute a regulatory network. The information that one can infer from its analysis provides a meaningful qualitative description about the dynamics of a GRN. Because of this, this approach has been used to study different biological models (88–90). In the Boolean network the state of each node is described by a discrete variable that takes the binary value: 0 (inhibited) or 1 (activated). In the present case, the nodes represent genes, nonetheless, they could also represent, proteins, RNA, morphogenes, etc. The links that join the nodes represent positive or negative regulations. The time evolution of the state of node *x*_*i*_, is determined by the Boolean function Ψ_*i*_ that depends on the states of its regulators *x*_*k*_, (with *k i*), located at other nodes as follows

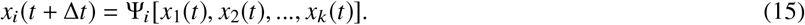

The Boolean function Ψ_*i*_ is discrete and follows the algebraic rules of Boole’s axiomatics. The stationary states of the dynamical mapping, called attractors, are determined by the condition *x*_*i*_(*t*) = *x*_*i*_(*t* + Δ*t*), for any Δ*t* > 0 that lead to activation or inhibition patterns defining a homeostatic state of the system. In the present context these attractor states represent normal, precancer or cancer cellular states.

An exhaustive exploration of the attractors of the breast cancer GRN was carried out using the BoolNet package of R (version 4.0.2). The knock-out (KO) and over-expression (OE) functions of each node were represented by a 0 (KO) or 1 (OE) value. The action of the altered node was also explored. In Figs. 11-12 are the tables that show all the identified system attractors. Many repeating attractors were found which were classified in groups and arranged in a single table that is presented in Fig. 11. These results are important because it is much simpler to analyze the GRN Boolean interactions and dynamics. These results can be used as a guide to built up and understand the complex dynamics of a continuous model. In the next section we use the properties and information of the Boolean network attractors to tune up some of the continuous model parameters values.

**Figure 11:**
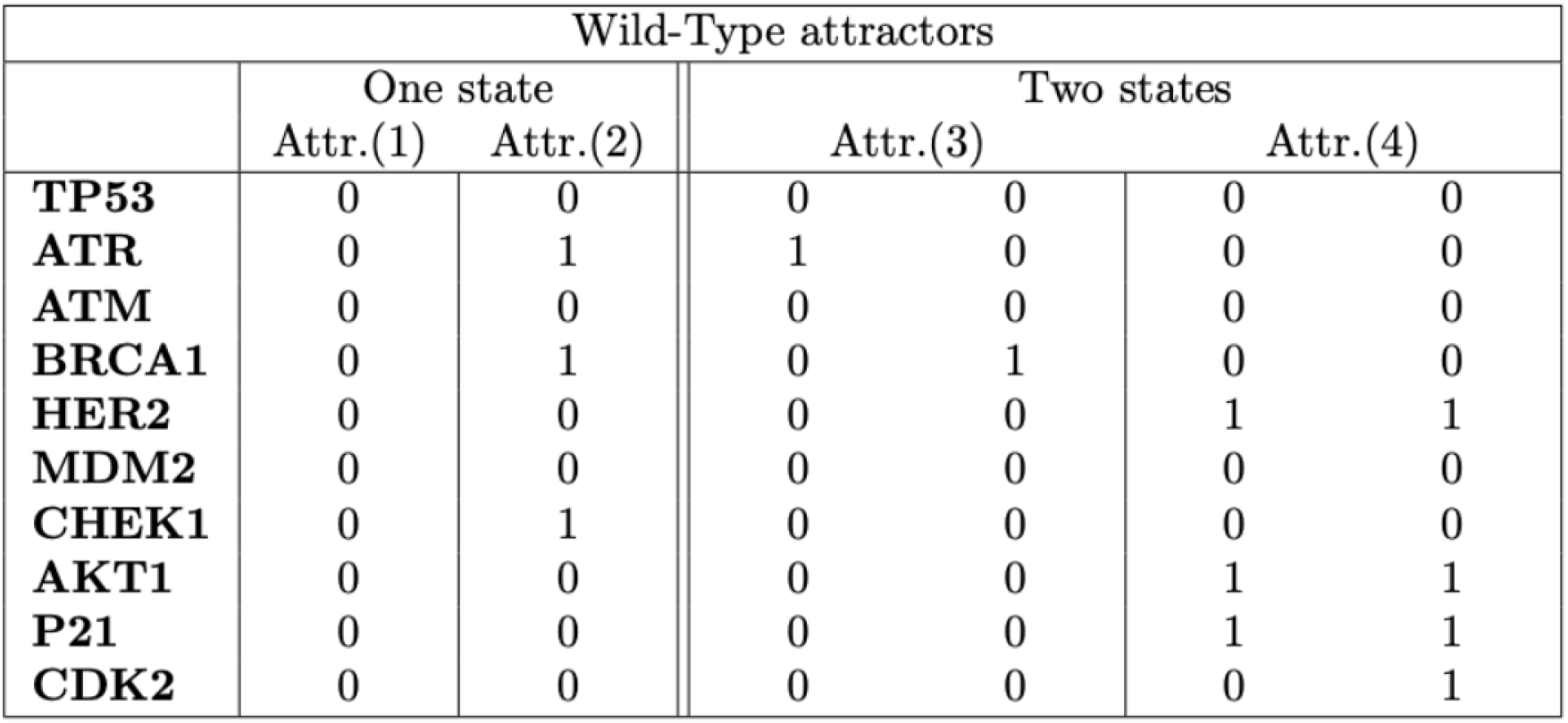
Wild Type attractors (Attr) obtained by using the Boolean representation of GRN.

All network attractors for breast cancer (see in Figs. 11-12) can be summarized as follows: a) The first attractor of Wild-Type character is identical for genes TP53-KO, ATR-KO, ATM-KO, BRCA-KO, HER2-KO, MDM2-KO, CHEK1-KO, AKT1-KO, P21-KO and CDK2-KO; b) The second attractor of Wild-Type character is identical for genes TP53-KO, ATR-OE, ATM-KO, BRCA1-OE, HER2-KO, MDM2-KO, MDM2-OE, CHEK1-OE, AKT1-KO, P21-KO and CDK2-KO; c) The third attractor of Wild-Type character is identical for genes TP53-KO, ATM-KO, HER2-KO, MDM2-KO, CHEK1-KO, AKT-KO, P21-KO and CDK2-KO; d) The fourth, and last attractor of Wild-Type character is identical for genes TP53-KO, ATR-KO, ATM-KO, BRCA1-KO, HER2-OE, MDM2-KO, CHEK1-KO, CHEK1-OE, AKT1-OE and P21-OE; e) Finally, the remaining attractors represent some other specific functions of some genes and are presented in the tables shown in Fig. 12.

**Figure 12:**
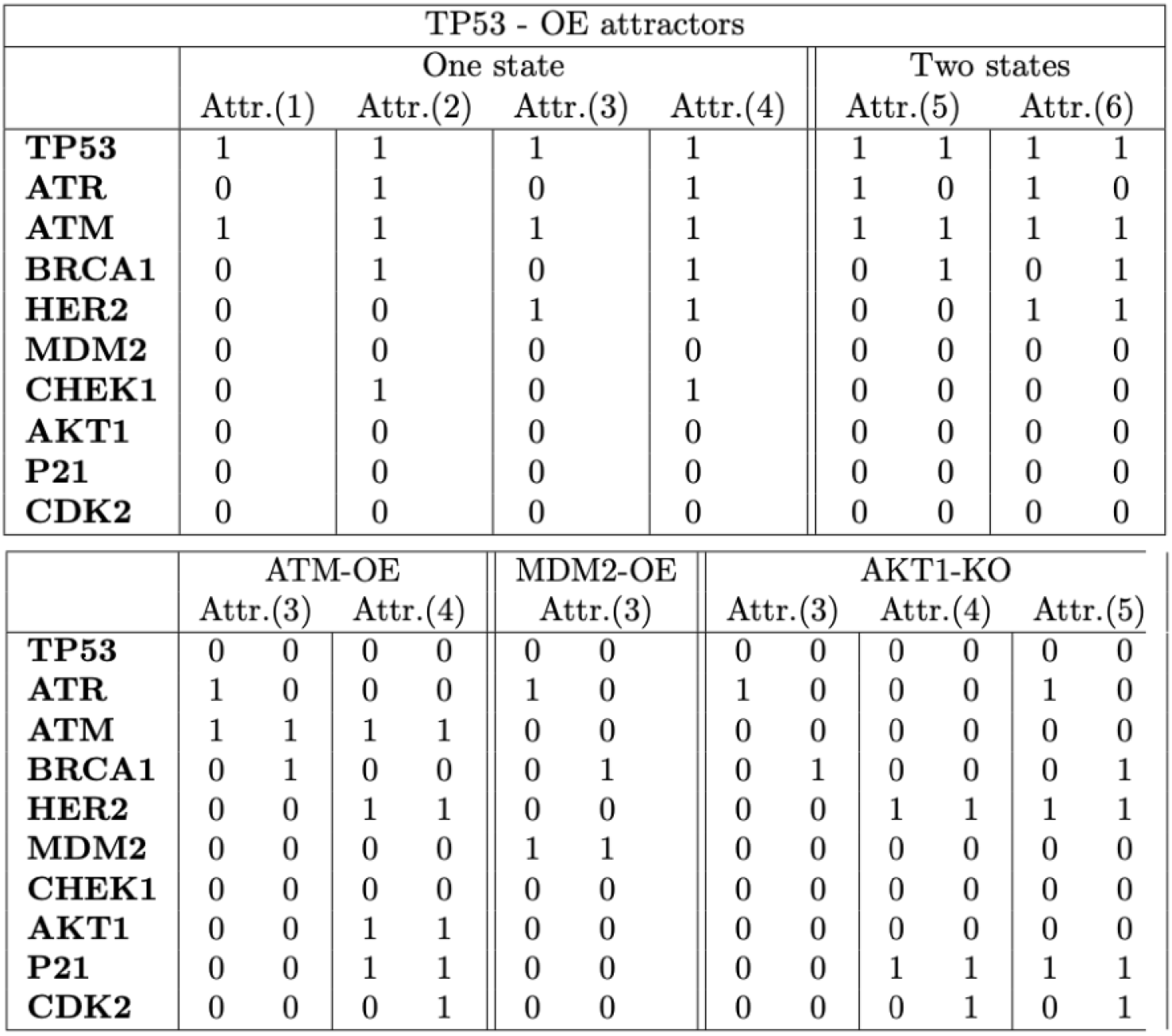
Breast cancer attractors (Attr) using over-expression (OE) and knock-out (KO) techniques on specific genes.

#### 1.2 Continuous model

Epigenetic mechanisms are essential for normal development and maintenance of tissue specific gene expression patterns. Disruption of these mechanisms may lead to altered cell functions. Because of this, epigenetics mechanisms have an important role in breast cancer development. It has been found that oxygen and estrogens located in the tumor microenvironment change the gene expression levels by activating or inhibiting genes which affect cell lineage (57). To model the effect of estrogens and oxygen in the the GRN dynamics we have used the gene interaction kinetics previously introduced in (86). -See Eqns (10) in the main text-. In this approach, the values of the state variables and parameters that control the dynamics of the GRN –shown in Fig. 1– vary continuously within a certain range as a result of the estrogens and oxygen concentrations gradients that lead to diverse genetic states. The temporal evolution of the state *x*_*i*_ of gene *i* is determined by its network kinetic interactions described by the following system of coupled ordinary differential equations:

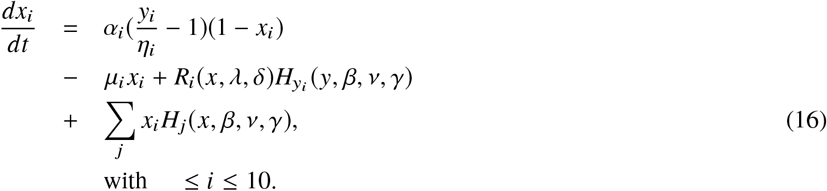

The parameters *α*_*i*_ represent the interaction rate constants between gene states and microenvironment. The action of the microenvironmental agents on gene activation is represented by *y*_*i*_ while *η*_*i*_ represents the activation-inhibition threshold parameter and *µ*_*i*_ is the self-degradation constant. The self-regulatory positive (+) and negative (−) feedback gene interactions are represented by the following equation,

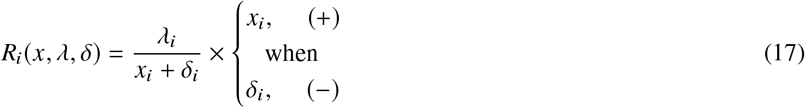

where *λ*_*i*_ is the activation constant and *δ*_*i*_ is the activation threshold. The genetic strength is represented by Hill function *H*_*j*_ as

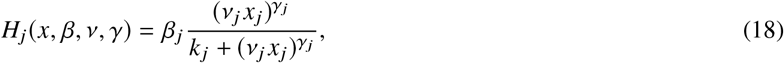

where β_*j*_ is the activation-inhibition parameters, *ν*_*j*_ is the action velocity at which gene *j* affects the state of gene *i*, the parameters *k* _*j*_ represent the sigmoidal function threshold, γ_*j*_ is the Hill coefficient which controls the steepness of the sigmoidal function and represents the cooperativeness of the regulatory transcription factor binding to genes. In the present case *k* _*j*_ = 1/2 for all genes, which means that all Hill functions activate/inhibit the corresponding gene state in the same manner. Moreover, the value of γ_*j*_ depends on the number of neighbor genes that contribute to the change of state of gene *j* as a cooperative system. In the Table (2) are presented the values of all the parameters involved in Eqns. (16,17,18). The values of these parameters were obtained from an exhaustive analysis using different algorithms (91–95) in order that the continuous dynamics yielded all gene states and attractors found in the Boolean model for wild -type, KO and OE states (see section 1.1).

**Table 2:**
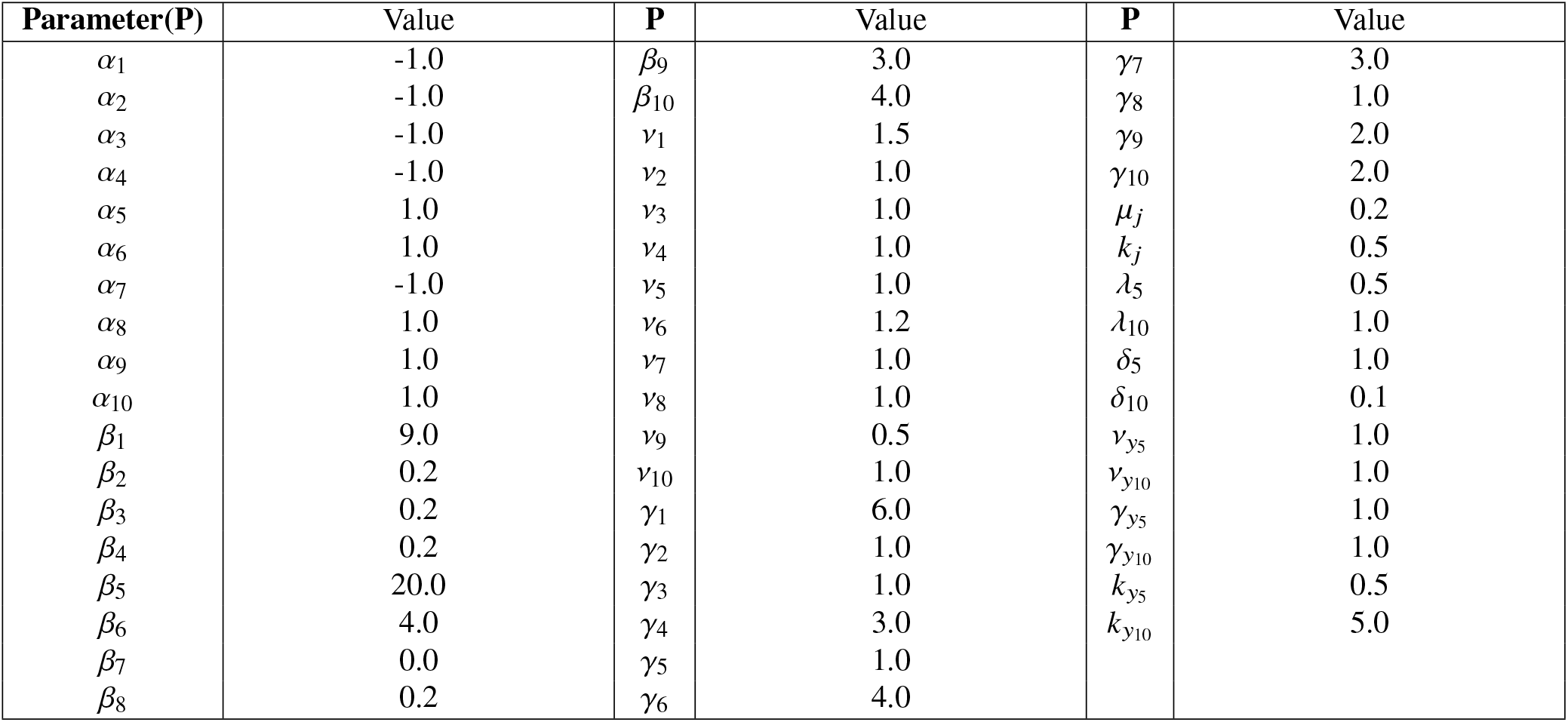
Parameter values for gene expression

| Parameter(P) | Value | P | Value | P | Value |
| --- | --- | --- | --- | --- | --- |
| $\alpha_1$ | -1.0 | $\beta_9$ | 3.0 | $\gamma_7$ | 3.0 |
| $\alpha_2$ | -1.0 | $\beta_{10}$ | 4.0 | $\gamma_8$ | 1.0 |
| $\alpha_3$ | -1.0 | $\nu_1$ | 1.5 | $\gamma_9$ | 2.0 |
| $\alpha_4$ | -1.0 | $\nu_2$ | 1.0 | $\gamma_{10}$ | 2.0 |
| $\alpha_5$ | 1.0 | $\nu_3$ | 1.0 | $\mu_j$ | 0.2 |
| $\alpha_6$ | 1.0 | $\nu_4$ | 1.0 | $k_j$ | 0.5 |
| $\alpha_7$ | -1.0 | $\nu_5$ | 1.0 | $\lambda_5$ | 0.5 |
| $\alpha_8$ | 1.0 | $\nu_6$ | 1.2 | $\lambda_{10}$ | 1.0 |
| $\alpha_9$ | 1.0 | $\nu_7$ | 1.0 | $\delta_5$ | 1.0 |
| $\alpha_{10}$ | 1.0 | $\nu_8$ | 1.0 | $\delta_{10}$ | 0.1 |
| $\beta_1$ | 9.0 | $\nu_9$ | 0.5 | $\nu_{y_5}$ | 1.0 |
| $\beta_2$ | 0.2 | $\nu_{10}$ | 1.0 | $\nu_{y_{10}}$ | 1.0 |
| $\beta_3$ | 0.2 | $\gamma_1$ | 6.0 | $\gamma_{y_5}$ | 1.0 |
| $\beta_4$ | 0.2 | $\gamma_2$ | 1.0 | $\gamma_{y_{10}}$ | 1.0 |
| $\beta_5$ | 20.0 | $\gamma_3$ | 1.0 | $k_{y_5}$ | 0.5 |
| $\beta_6$ | 4.0 | $\gamma_4$ | 3.0 | $k_{y_{10}}$ | 5.0 |
| $\beta_7$ | 0.0 | $\gamma_5$ | 1.0 | | |
| $\beta_8$ | 0.2 | $\gamma_6$ | 4.0 | | |

In Table (3) one identifies the GRN Boolean attractors with those corresponding to the continuous representation by assigning the appropriate genes activation/inhibition threshold parameter values. A detailed and exhaustive analysis of these parameters led to the conclusion that the threshold activation values should be equal to 0.8. This result is in full agreement with most of the thresholds activation values found in Ref. (86), where a stochastic model was analyzed and the thresholds were expressed as percentages. In all microarrays obtained with the continuous model genes expression for precancer genotype where found in the parameter range (0, 0.8). On the other hand, genes inhibition were obtained with a threshold equal to zero whereas genes activations were obtained for values higher than the threshold values. The set of genetic expressions in table **??** define the corresponding cellular phenotypes which are: (i) normal, (ii) precancerous, and (iii) cancer or malignant.

**Table 3:** Gene expression value in each state

| Gene | Boolean model |  | Continuous model |  |  |
| --- | --- | --- | --- | --- | --- |
|  | Cancer | Normal | Cancer | Precancer | Normal |
| TP53(x1) | 0 | 1 | 0 | $<0.8 \leq$ | 1 |
| ATR(x2) | 0 | 1 | 0 | $<0.8 \leq$ | 1 |
| ATM(x3) | 0 | 1 | 0 | $<0.8 \leq$ | 1 |
| BRAC1(x4) | 0 | 1 | 0 | $<0.8 \leq$ | 1 |
| HER2(x5) | 1 | 0 | 1 | $\geq 0.8 >$ | 0 |
| MDM2(x6) | 1 | 0 | 1 | $\geq 0.8 >$ | 0 |
| CHEK1(x7) | 0 | 1 | 0 | $<0.8 \leq$ | 1 |
| AKT1(x8) | 1 | 0 | 1 | $\geq 0.8 >$ | 0 |
| P21(x9) | 1 | 0 | 1 | $\geq 0.8 >$ | 0 |
| CDK2(x10) | 1 | 0 | 1 | $\geq 0.8 >$ | 0 |

Figure 13 shows a microarray that represents the attractors obtained from the dynamics of the GRN continuous model shown in Fig. 1 in the main text. To reproduce the GRN Boolean attractors shown in Figs. 11 and 12, an exhaustive and sensitive parameter analysis was performed to obtain the initial threshold values parameters *η*_*oi*_, that led to the representation of precancer phenotypes indicated in the microarrays shown in Fig. 13. Attractor states **a** and **c** are identified with normal phenotypes, whereas attractor states **b, d, e, f, g, h**, and **i** can be identified with a precancer phenotype since some genes are either over-expressed or inhibited permanently in the Boolean GRN representation (see Figs. 11 and 12). Finally, the state **j** is the only attractor that one can identify with cancer phenotype.

**Figure 13:**
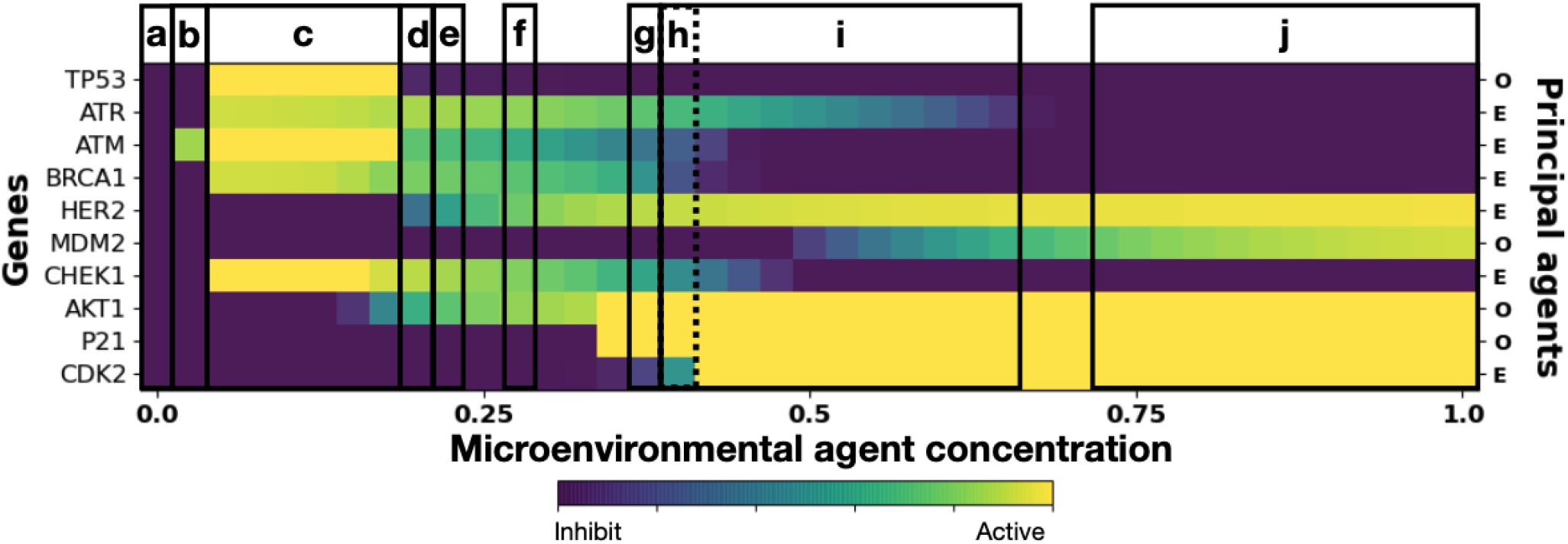
Microarrays showing genes expression and cell phenotypes. Normal cell phenotypes are represented by the states **a** and **c**. The states **b, d, e, f, g, h**, and **i** are identified with attractors in which some genes are Over-Expressed or Knocked-Out, suggesting a precancer phenotype. The state **j** is identified with the attractor that represents cancer phenotypes. The initial threshold parameters for the set of ten genes are: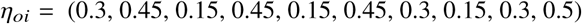.

Fig. 14 present a microarray that indicates the effect of increasing estrogens threshold values on the expression of all genes. The initial threshold values η_*oi*_ were increased in the proportions indicated in the figure caption. The microarray presented in Fig. 15 was obtained by increasing the estrogens threshold value in gene HER2. The initial threshold value η_*oi*_ was increased in the proportions indicated in the figure caption. By comparing these microarrays it is found that the precancer phenotype decreases more when an increase in estrogens threshold value occurs in gene HER2 rather than in all other genes. It has been found that overexpression of HER2 leads to malignant phenotype and accelerates tumorigenesis (24). This results reaffirms the role of estrogen concentration in changing the gene expression levels and as a result the occurrence of different phenotypes. This analysis is in complete agreement with previous findings (17, 18, 20, 56, 70) and give support to the model and results presented in this paper.

**Figure 14:**
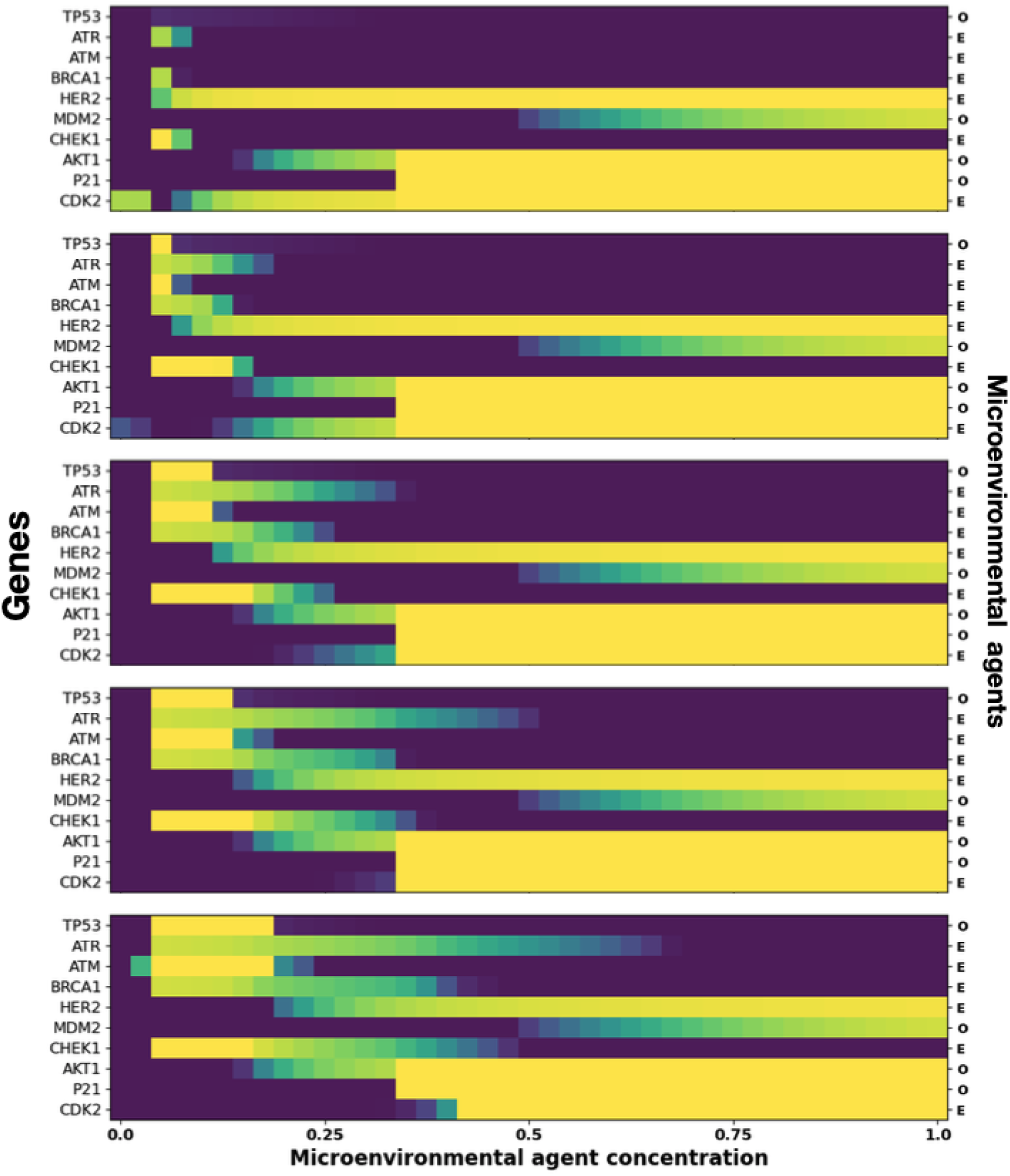
Microarrays showing the effects of increasing estrogens threshold values in the expression of all ten genes. From top to bottom, the increase occurs as follows: 10%, 25%, 50%, 75% and 100%, as compared to the values assigned to obtain the microarrays shown in Fig. 13. The parameters and color bar are the same as in Fig. 13

**Figure 15:**
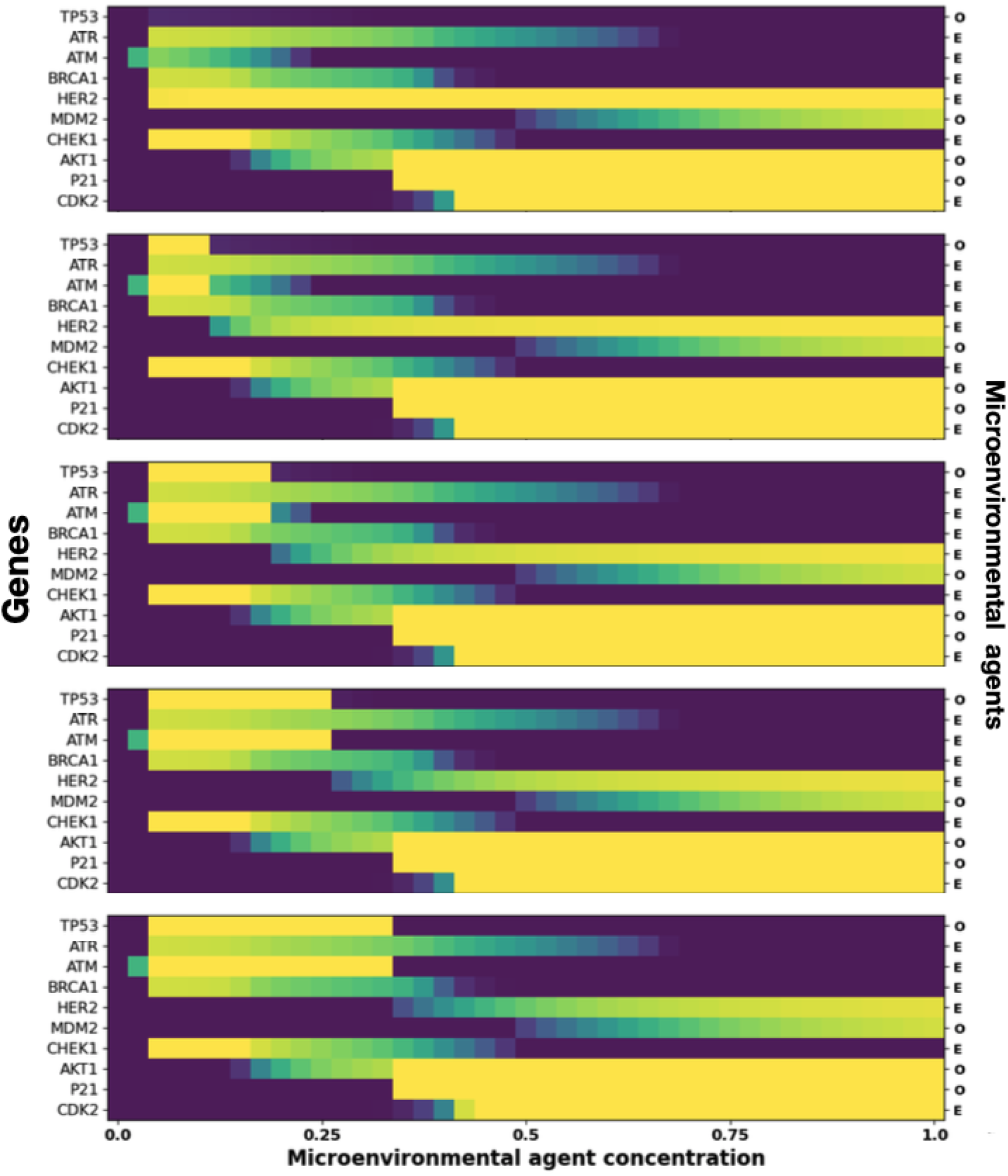
Microarrays showing the effects of increasing estrogens threshold value in gene HER2. From top to bottom, the increase is as follows: 0.1%, 50%, 100%, 150% and 200% as compared to the values assigned in Fig. 13. The parameters and color bar are the same as in Fig. 13

